# An enteric neuron-expressed variant ionotropic receptor detects ingested salts to regulate salt stress resistance

**DOI:** 10.1101/2025.04.11.648259

**Authors:** Jihye Yeon, Laurie Chen, Nikhila Krishnan, Sam Bates, Charmi Porwal, Piali Sengupta

**Author notes:** Corresponding authors: P.S.; J.Y.

## Abstract

The detection of internal chemicals by interoceptive chemosensory pathways is critical for regulating metabolism and physiology. The molecular identities of interoceptors, and the functional consequences of chemosensation by specific interoceptive neurons remain to be fully described. The *C. elegans* pharyngeal neuronal network is anatomically and functionally homologous to the mammalian enteric nervous system. Here, we show that the I3 pharyngeal enteric neuron responds to cations via an I3-specific variant ionotropic receptor (IR) to regulate salt stress tolerance. The GLR-9 IR, located at the gut lumen-exposed sensory end of I3, is necessary and sufficient for salt sensation, establishing a chemosensory function for IRs beyond insects. Salt detection by I3 protects specifically against high salt stress, as *glr-9* mutants show reduced tolerance of hypertonic salt but not sugar solutions, with or without prior acclimation. While cholinergic signaling from I3 promotes tolerance to acute high salt stress, peptidergic signaling from I3 during acclimation is essential for resistance to a subsequent high salt challenge. Transcriptomic analyses show that I3 regulates salt tolerance in part via regulating the expression of osmotic stress response genes in distal tissues. Our results describe the mechanisms by which chemosensation mediated by a defined enteric neuron regulates physiological homeostasis in response to a specific abiotic stress.

## INTRODUCTION

While foraging for food, animals often encounter toxic substances that adversely affect health and longevity. Consequently, animals have evolved multiple mechanisms to identify and cope with harmful chemicals. A first line of defense involves behavioral avoidance mediated by exteroceptive chemosensory neurons present in olfactory and taste organs (Ferkey et al., 2021; Taruno et al., 2021). If ingested, chemicals in the gastrointestinal tract are detected directly or indirectly by non-neuronal cells, as well as gut-intrinsic enteric neurons and extrinsic sensory afferents, to regulate and maintain metabolic homeostasis (Furness et al., 1999; Miguel-Aliaga et al., 2018; Prescott and Liberles, 2022; Spencer and Hu, 2020). In contrast to the well-described exteroceptive chemotransduction pathways, the molecular mechanisms and functional roles of interoceptive chemosensation remain to be fully described.

Inorganic cations such as Na^+^ and Ca^2+^ and related salts are essential for the regulation of a range of physiological processes and are typically attractive at low concentrations (Berridge et al., 2003; Taruno and Gordon, 2023). However, high salt concentrations are harmful (Hunter et al., 2022; Rucker et al., 2018), and are detected by gustatory neurons to drive rejection and avoidance, as well as by interoceptive neurons in multiple organs including the gut and brain to modulate salt and fluid intake (Stocker et al., 2024; Taruno and Gordon, 2023). Although the toxic effects of high salt are generally attributed to osmotic stress, comparisons with cellular responses elicited by equiosmolar sugar solutions suggest that individual hypertonic solutes can be discriminated and their concentrations monitored to regulate different physiological responses (Kinsman et al., 2017; Lamitina et al., 2004; Stocker et al., 2024).

Salt sensation is mediated by a diverse range of molecules across species. These include amiloride-sensitive Na^+^ channels in mammals, and an expanded family of variant ionotropic receptors (IRs) in *Drosophila* (Stocker et al., 2024; Taruno and Gordon, 2023). In *Drosophila*, distinct IR protein complexes expressed in different external and internal taste sensilla have been implicated in detecting a range of salt concentrations (Kim et al., 2024; Sang et al., 2024; Taruno and Gordon, 2023). Although IR proteins are encoded by the genomes of all protostomes (Croset et al., 2010), chemosensory roles for these proteins in species beyond insects remain to be definitively established.

The nematode *C. elegans* is also attracted to low salt but avoids high salt using a small set of ciliated chemosensory neurons present in sense organs of the head and tail (Ferkey et al., 2021). Exposure to high NaCl rapidly paralyzes and eventually kills worms but prolonged exposure to sub-lethal NaCl concentrations increases tolerance upon subsequent high NaCl exposure (acclimation) (Lamitina et al., 2004). Worms are also paralyzed rapidly in high sucrose concentrations (Lamitina et al., 2004), implying that the effects of high salt are due to hypertonic stress. However, worms survive to different extents on equiosmolar NaCl and sugar suggesting that as in mammals, unidentified salt-specific sensors and pathways may regulate salt stress (Lamitina et al., 2004).

Food ingestion in *C. elegans* is primarily driven by the foregut or pharynx (Albertson and Thomson, 1976; Avery and You, 2012). This neuromuscular organ is innervated by a densely interconnected, autonomously functioning network of 20 pharyngeal neurons that exhibit developmental, anatomical, and functional homology to the mammalian enteric nervous system (Albertson and Thomson, 1976; Cook et al., 2020; Vidal et al., 2022). 15/20 of these pharyngeal neurons (henceforth referred to as pharyngeal enteric neurons; PENs) exhibit sensory features, although to date, only two PENs have been shown to respond to bacterial compounds to regulate feeding and foraging (Bhatla and Horvitz, 2015; Rhoades et al., 2019).

Here we show that the single I3 PEN in *C. elegans* responds to a range of monovalent and divalent cations and regulates salt stress resistance. Salt detection is mediated by the I3-specific GLR-9 IR-related protein which is both necessary and sufficient for salt responses. Salt sensation by I3 is transmitted via cholinergic and peptidergic signaling to increase tolerance to high salt but not high sugars in unacclimated and acclimated animals, respectively. Salt acts via both GLR-9-dependent and -independent pathways to alter the expression of genes in distal tissues including the epidermis and gut that enable salt stress resistance; misregulation of this gene expression program in *glr-9* mutants likely underlies their salt-sensitivity phenotype. Together, our results identify a chemosensory function of an IR in nematodes, and elucidate how a single enteric neuron detects and relays the presence of ingested cations to counter salt toxicity.

## RESULTS

### The GLR-9 tuning and GLR-7 co-receptor IRs are co-localized to the gut lumen-exposed sensory ending of the I3 pharyngeal enteric neuron

The *C. elegans* genome encodes 15 predicted ionotropic receptors related to glutamate-gated cation channels (Brockie et al., 2001; Hobert, 2013) (Fig. 1a). Of these, seven exhibit features characteristic of variant sensory IRs in insects including amino terminal domains of varying lengths, the lack of canonical glutamate-interacting residues in their ligand binding domains, and loss of highly conserved sequences essential for regulating ion permeability in their pore-forming domains (Benton et al., 2009; Brockie et al., 2001; Croset et al., 2010) (Fig. 1a, Fig. S1a). Examination of the neuronal transcriptome database (Taylor et al., 2021) showed that six of these IR-like genes are exclusively or highly expressed in subsets of PENs (Brockie et al., 2001) (Fig. S1b). In particular, the *W02A2.5* IR gene (henceforth referred to as *glr-9*) is expressed only in the single I3 PEN (Fig. S1b), suggesting that *glr-9* may play a specialized sensory role in this neuron.

**Fig. 1.**
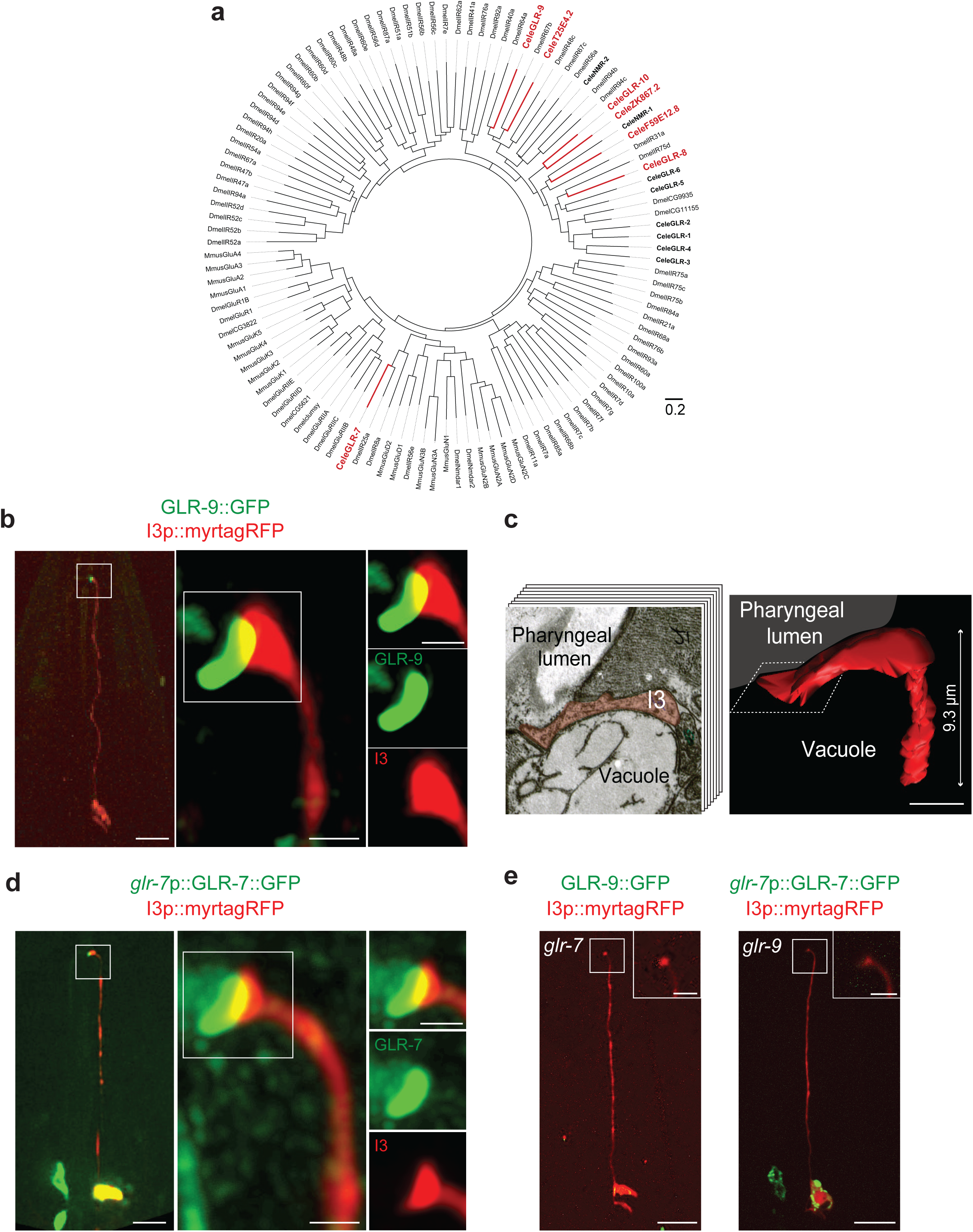
GLR-9 and GLR-7 are localized at the pharyngeal lumen-exposed sensory ending of I3. **a)** A rooted polar phylogenetic tree of ionotropic glutamate receptor sequences from *Drosophila* (Dmel), *C. elegans* (Cele; bolded) and mouse (Mmus). Variant *C. elegans* IRs are indicated in bold red; canonical ionotropic glutamate receptors are in bold black. The scale bar represents the branch length, with each unit equivalent to 0.2 nucleotide substitutions per site. **b,d)** Representative images of GLR-9 (b) or GLR-7 (d) fusion protein localization in I3. The I3 sensory ending is boxed and magnified in images on the right. GLR-9 is tagged with GFP at its endogenous locus; *glr-7*p::GLR-7::GFP is expressed from an extrachromosomal array. I3 is marked via the expression of *glr-9*p::myrtagRFP. Anterior is at top in all images. Scale bars: 10 μm (images at left); 0.5 μm (images at center and right). **c)** (Left) Cross-section electron microscope image approximately 20 μm from the worm nose showing the pharyngeal lumen, a vacuole, and I3 (shaded). (Right) 3D reconstruction of the I3 distal ending from electron microscope serial sections. The location of the pharyngeal lumen is shown. The length of the reconstructed I3 ending is indicated at right. A dashed box indicates the approximate position of the electron microscope cross-section image shown at left. Arrowhead indicates the sensory ending. Scale bar: 0.25 μm. **e)** Localization of GLR-9::GFP or GLR-7::GFP in I3 in *glr-7(tm2877)* or *glr-9(oy180)* mutant backgrounds, respectively. Anterior is at top. The I3 sensory ending is boxed and magnified in the insets. Scale bars: 10 μm; 2.5 μm (insets).

Consistent with transcriptomics data, an endogenous GFP-tagged GLR-9 fusion protein was expressed only in I3 (Fig. 1b). The GLR-9 fusion protein localized to a structure that is present asymmetrically at the I3 ending and protrudes from the cell surface towards the interior of the animal (Fig. 1b, Movie S1). Reconstruction of the sensory ending of I3 from serial section electron micrographs also showed a structure that is expanded and curved (Fig. 1c, Movie S2) (Albertson and Thomson, 1976; Cook et al., 2020), and that may represent the domain occupied by GLR-9. The internal side of this ending is directly adjacent to and is likely exposed to the pharyngeal lumen (Fig. 1c, Movie S2). However, I3 is not ciliated and does not express ciliary genes (Taylor et al., 2021), leaving open the question of the identity of this structure. We infer that GLR-9 may be positioned in I3 to directly sample ingested molecules present in the pharyngeal lumen.

Sensory IRs in insects typically function as heterotetramers comprised of a stimulus-specific tuning receptor and one or more co-receptor(s) that traffic the IR complex to the ciliated endings of sensory neurons (Abuin et al., 2011; Abuin et al., 2019; Ai et al., 2013). Similar to other tuning IRs, GLR-9 does not contain an extended amino-terminal domain (Fig. S1a), suggesting that this IR may also act as a tuning receptor. Among the characterized IR co-receptors, IR25a is deeply conserved in all protostomes and is considered the ancestral chemosensory IR (Croset et al., 2010). The *glr-7* IR25a ortholog in *C. elegans* (Croset et al., 2010) (Fig. 1a) is expressed in a small number of pharyngeal and non-pharyngeal neurons with the highest expression in I3 (Fig. S1b) (Taylor et al., 2021). A GFP-tagged GLR-7 fusion protein expressed in I3 was also localized to the sensory ending of I3 (Fig. 1d). Localization of GLR-9 was abolished in *glr-7* mutants and *vice versa* (Fig. 1e). Together, these results suggest that GLR-9 and GLR-7 may comprise a sensory receptor complex in the I3 PEN.

### GLR-9/GLR-7 are necessary and sufficient to mediate I3 responses to high concentrations of monovalent and divalent cations

To determine whether the GLR-9/GLR-7 complex detects chemicals present in the pharyngeal lumen, we surveyed chemosensory stimuli-evoked intracellular calcium dynamics in I3 neurons expressing the GCaMP6s calcium sensor. I3 responded robustly to both bacterial supernatant as well as bacterial growth media (LB: Lysogeny or Luria Broth) alone (Fig. S2a) suggesting that I3 responds to one or more chemical components in LB. Further analyses showed that I3 responded to a range of monovalent and divalent salts including NaCl, NaAc, KPO_4_, NH_4_Cl, MgSO_4_, NH_4_Ac, KCl, NaPO_4_, (NH_4_)_2_SO_4_, and CaCl_2_ (Fig. 2a, Fig. S2b). Replacement of Na^+^ with N-methyl-D-glucamine (NMDG) in NMDG-Cl^-^ failed to elicit a response, whereas replacing Cl^-^ with gluconate in Na^+^-gluconate resulted in increased intracellular calcium levels (Fig. 2b, Fig. S2c), suggesting that I3 responds non-selectively to cations. No responses were observed to equiosmolar concentrations of fructose, sorbitol, or glycerol (Fig. 2c, Fig. S2d), indicating that I3 does not respond to high osmolarity.

**Fig. 2.**
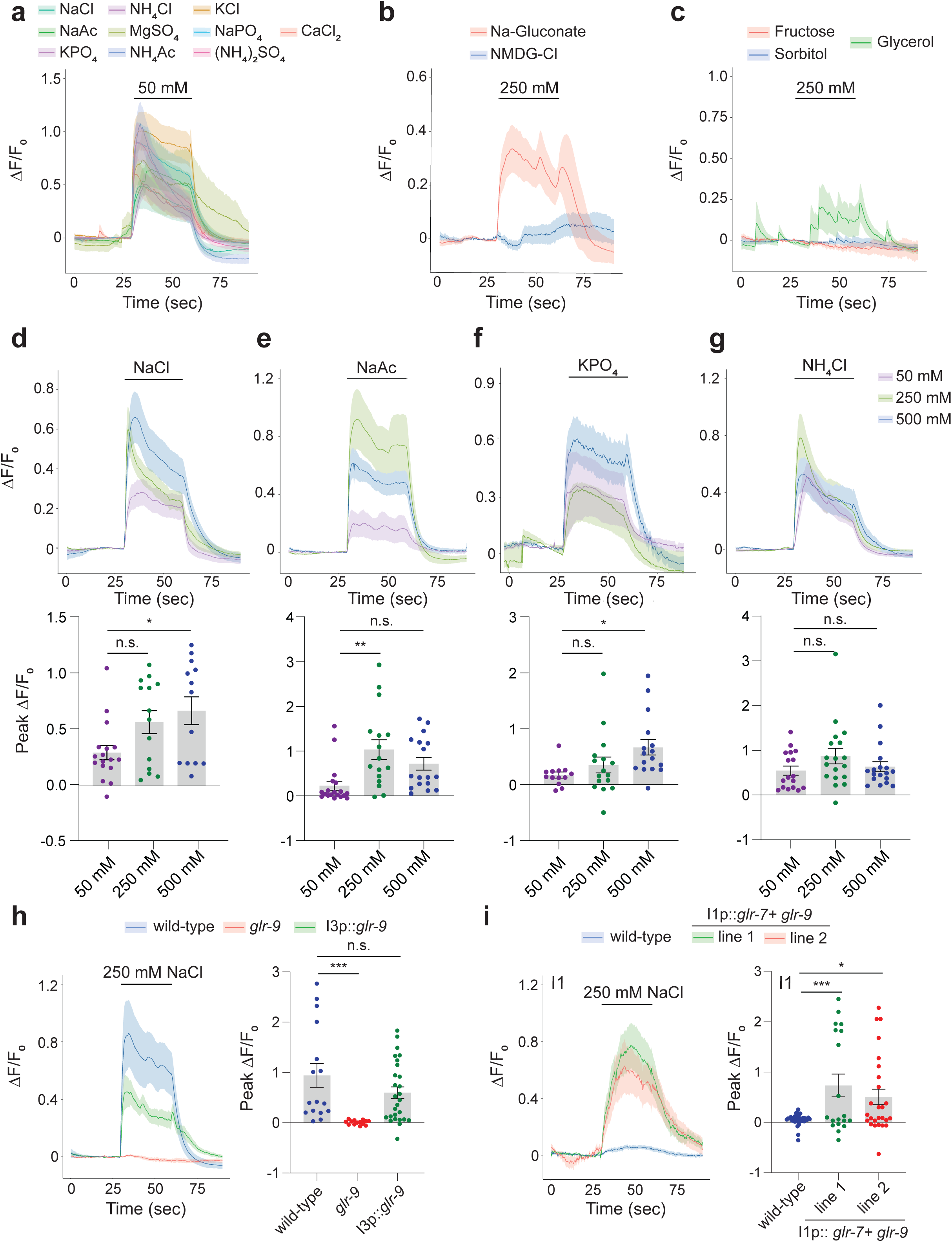
I3 responds to multiple monovalent and divalent cations in a GLR-9-dependent manner. **a-c)** Average changes in GCaMP6s fluorescence in I3 in response to a 30 sec pulse of the indicated salts (a,b) or sugars (c) at the shown concentrations. Quantification of peak fluorescence intensities are shown in Fig. S2b-d. **d-g)** Average (top) and peak intensity changes (bottom) of GCaMP6s fluorescence in I3 in response to a 30 sec pulse of different concentrations of NaCl, NaAc, KPO_4_, and NH_4_Cl. * and ** indicate different from corresponding 50 mM values at P< 0.05 and 0.01, respectively (one-way ANOVA and Dunnett’s test). **h)** Average (left) and peak intensity changes (right) of GCaMP6s fluorescence in I3 in response to a 30 sec pulse of 250 mM NaCl in animals of the indicated genotypes. The *glr-9(oy180)* allele was used. Wild-type *glr-9* sequences were expressed in I3 under the *glr-9* promoter. Rescue data shown are combined values from two independent transgenic lines; responses of each line were similar. *** indicates different from wild-type at P<0.001 (one-way ANOVA and Dunnett’s test). **i)** Average (left) and peak intensity changes (right) of GCaMP6s fluorescence in wild-type I1 neurons or in I1 misexpressing GLR-7 and GLR-9 to a 30 sec pulse of 250 mM NaCl. Expression in I1 was driven under the *lgc-8* promoter. Responses from two independent misexpressing lines are shown. * and *** indicate different from wild-type I1 at P<0.01 and 0.001, respectively (one-way ANOVA and Dunnett’s test). Shaded regions in all traces indicate SEM. Each dot in the scatter plots is the value from a single neuron. Horizontal lines in scatter plots indicate the mean; errors are SEM. Data shown are from 2-3 independent experiments each. n.s. – not significant.

I3 responded largely in a dose-dependent manner to NaCl, NaAc, KPO_4_ and NH_4_Cl (Fig. 2d-g). Responses at higher salt concentrations appeared to be tonic with little to no desensitization, and returned to baseline upon removal of the stimulus (Fig. 2d-h, Fig. S2e). Salt responses were maintained in *unc-13* and *unc-31* mutants which abolish synaptic or neuropeptidergic transmission, respectively (Sieburth et al., 2007; Speese et al., 2007) (Fig. S2f), suggesting that I3 responds to salts cell-autonomously, although we are unable to exclude communication via gap junctions. Salt responses in I3 were also unaltered in animals mutant for the *tax-4* cyclic nucleotide-gated and *osm-9* TRPV channel genes required broadly for chemosensation in the ciliated chemosensory neurons of the head amphid sense organs, as well as in animals mutant for the *che-1* terminal selector transcription factor that specifies the fate of the salt-sensing ASE amphid chemosensory neurons (Fig. S2g) (Etchberger et al., 2007; Ferkey et al., 2021; Uchida et al., 2003). All examined salt responses in I3 were abolished in *glr-9* mutants; responses were restored in *glr-9* mutants upon I3-specific expression of wild-type *glr-9* sequences (Fig. 2h, Fig. S2h). We conclude that GLR-9/GLR-7 mediate I3 responses to a broad range of salts.

We next tested whether GLR-9/GLR-7 are also sufficient to confer salt responses onto a non salt-sensing neuron. Misexpression of GFP-tagged GLR-9 with or without GLR-7 in the salt-sensing ASE chemosensory neurons resulted in protein accumulation in the soma and failure to traffic into sensory cilia (Fig. S2i). Reasoning that pathways required for trafficking and localization of sensory IRs may be conserved in PENs, we misexpressed GLR-9 and GLR-7 in the I1 PENs whose distal non-ciliated sensory endings are also apposed to the gut lumen (Albertson and Thomson, 1976; Cook et al., 2020). Co-expression of both proteins but not each protein alone led to localization to the I1 sensory endings similar to the pattern in I3 (Fig. S2j). Although I1 did not respond to NaCl, misexpression of both GLR-9 and GLR-7 in I1 was sufficient to confer responses to NaCl (Fig. 2i). These observations indicate that GLR-9 and GLR-7 are both necessary and sufficient to confer salt responses in PENs.

### GLR-9-mediated salt sensation promotes salt stress resistance

To examine the functional consequences of I3-mediated salt sensation, we assessed multiple salt-regulated behaviors in *glr-9* mutants. Salt chemotaxis behaviors were examined in microfluidics behavioral arenas which allow analyses of behavioral responses to the same salt concentrations as those used to examine neuronal calcium responses (Albrecht and Bargmann, 2011; Khan et al., 2022). Wild-type animals were robustly attracted to a central stripe of 75 mM NaCl but avoided 250 mM NaCl (Fig. S3a). *glr-9* mutants exhibited no defects in either salt attraction or avoidance (Fig. S3a), and also exhibited robust avoidance of high glycerol concentrations (Fig. S3b). *glr-9* mutants also exhibited no defects in pharyngeal pumping frequency in low and high salt concentrations in the presence of bacterial food (Fig. S3c). These observations suggest that salt-sensing by I3 may contribute to organismal responses other than taxis or feeding.

*C. elegans* is typically grown on media containing 50 mM NaCl which is considered isotonic. When moved to media containing >500 mM NaCl, animals rapidly lose cell volume, become immotile due to loss of turgor pressure, and eventually die (Choe and Strange, 2007a; Lamitina et al., 2004). Since salt-specific pathways independent of hypertonicity have been proposed to also regulate salt toxicity (Lamitina et al., 2004), we tested the notion that I3-mediated salt sensation regulates high salt tolerance. Wild-type animals became rapidly immotile when transferred from 50 mM NaCl to 500 mM NaCl or high concentrations of other I3-sensed salts including NaAc, NH_4_Cl, or KPO_4_ (Fig. 3a). *glr-9* mutants were consistently paralyzed more rapidly than wild-type animals when moved from low to high NaCl (Fig. 3b), suggesting decreased salt tolerance. However, *glr-9* mutants did not exhibit enhanced sensitivity to equiosmolar sucrose solutions, (Fig. 3c, Fig. S3d) further highlighting the salt-specificity of I3 responses. We conclude that I3 activation specifically by salt is partly protective against acute high salt stress.

**Fig. 3.**
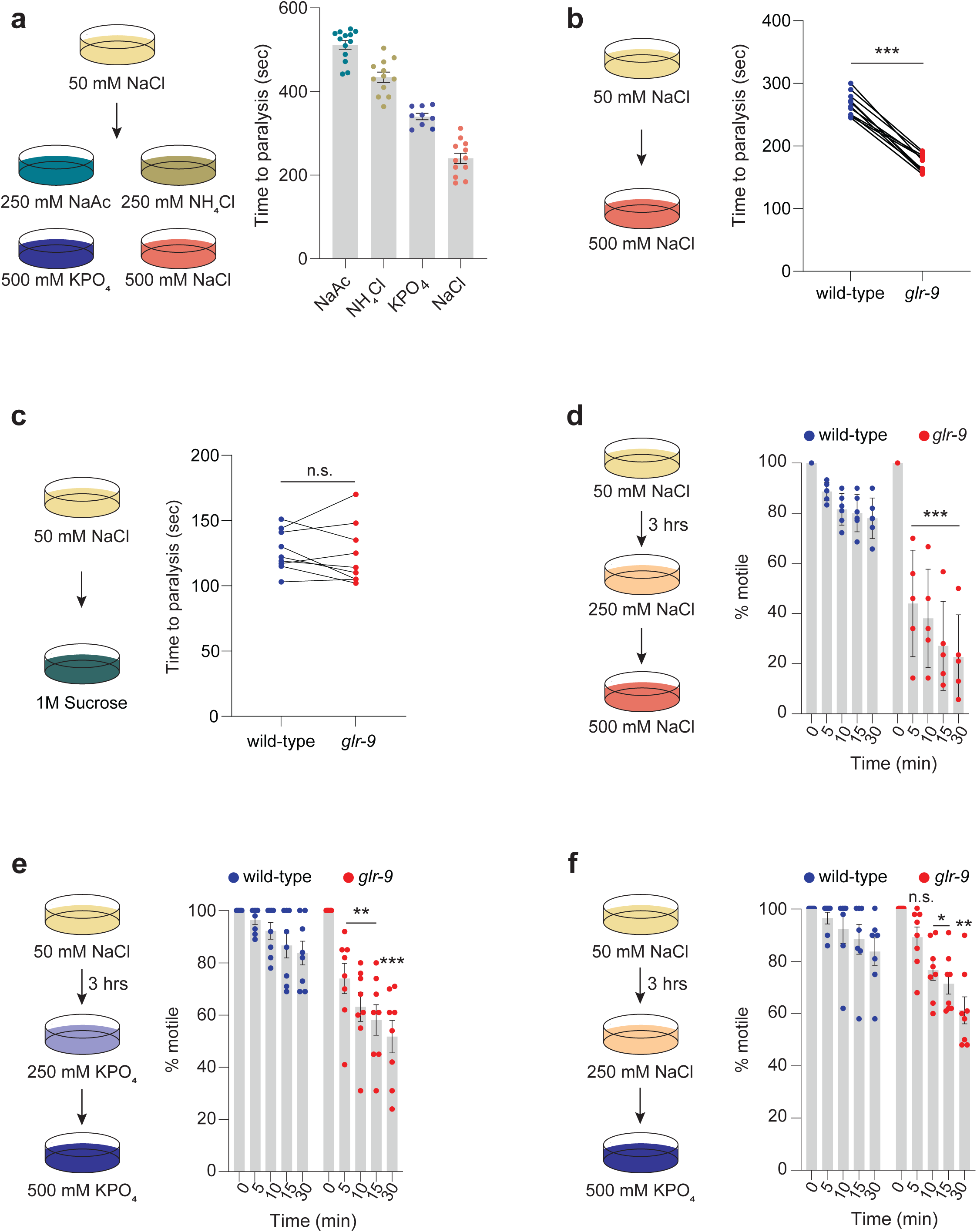
*glr-9* mutants exhibit reduced tolerance of high salt. **a)** (Left) Cartoon of experimental conditions. (Right) Quantification of time to paralysis upon shifting animals to each salt at concentrations indicated in the cartoon. Each dot is the time at which all ten animals in a single assay are immotile following the shift to high salt. **b,c)** Quantification of time to paralysis upon shifting animals of the indicated genotypes from 50 to 500 mM NaCl or equiosmolar 1 M sucrose (cartoon at left). Each dot is the time at which all ten animals in a single assay are immotile. Wild-type and *glr-9* mutants were examined in parallel in the same assay. *** indicates different from wild-type at P<0.001 (t-test). **d-f)** Percentage of animals of the indicated genotypes that are motile at each time point following a shift from the acclimation plate (250 mM salt for 3 hrs) to 500 mM salt (cartoons at left). Each dot is the value from a single assay; n=30 animals per assay. *, **, and *** indicate different from wild-type at the corresponding time at P<0.05, 0.01, and 0.001 (one-day ANOVA and Dunnett’s test). The *glr-9(oy180)* allele was used in all experiments. Horizontal lines in scatter plots indicate the mean; errors are SEM. Data shown are from 2-3 independent experiments each. n.s. – not significant.

While animals are highly sensitive to acute salt stress, pre-adaptation to 200–250 mM NaCl for 24 hrs allows *C. elegans* to tolerate subsequent exposure to >500 mM NaCl (Lamitina et al., 2004; Urso et al., 2020). This acclimation is thought to be mediated in part via increased expression of enzymes required for the production of intracellular osmolytes which enable osmotolerance (Choe, 2013; Lamitina et al., 2004; Urso and Lamitina, 2021). We examined whether *glr-9* mutants exhibit reduced salt tolerance even upon acclimation. Wild-type animals pre-exposed to 250 mM NaCl or 250 mM KPO_4_ for as little as 3 hrs continued to exhibit normal locomotion and pharyngeal pumping for >30 mins when placed on 500 mM NaCl or 500 mM KPO_4_, respectively (Fig. 3d-e). However, a significant fraction of *glr-9* mutants again became immotile within 5-10 mins when placed on high concentrations of either NaCl or KPO_4_ (Fig. 3d-e). Acclimation was unaffected in animals mutant for signaling genes required for sensory transduction in ciliated chemosensory neurons (Fig. S3e), indicating that these neurons do not contribute to high salt resistance. Since I3 exhibits responses to multiple salts, we next tested whether acclimation to sub-lethal concentrations of one salt enables tolerance of high concentrations of a different salt. Pre-exposure to 250 mM NaCl was sufficient to partly increase tolerance of wild-type but not *glr-9* mutant animals to 500 mM KPO_4_ (Fig. 3f). Taken together, we conclude that GLR-9-dependent salt responses in I3 are necessary for tolerance of high salts in both unacclimated and acclimated conditions.

### Cholinergic and FLP-6 neuropeptide-mediated signaling from I3 are necessary for high salt tolerance under different conditions

Osmoregulatory mechanisms operate in multiple tissues and organs to regulate osmotolerance in *C. elegans* (Choe, 2013; Urso and Lamitina, 2021), suggesting that I3 may signal to one or more of these tissues to regulate salt stress resistance. I3 is a cholinergic neuron and also expresses multiple neuropeptides (Cook et al., 2020; Taylor et al., 2021). Knocking out the *unc-17* vesicular acetylcholine transporter specifically in I3 using Cre-Lox mediated recombination (Huang et al., 2023) resulted in rapid paralysis upon exposure to high salt similar to *glr-9* mutants (Fig. 4a), indicating that cholinergic signaling from I3 is necessary for acute salt tolerance.

**Fig. 4.**
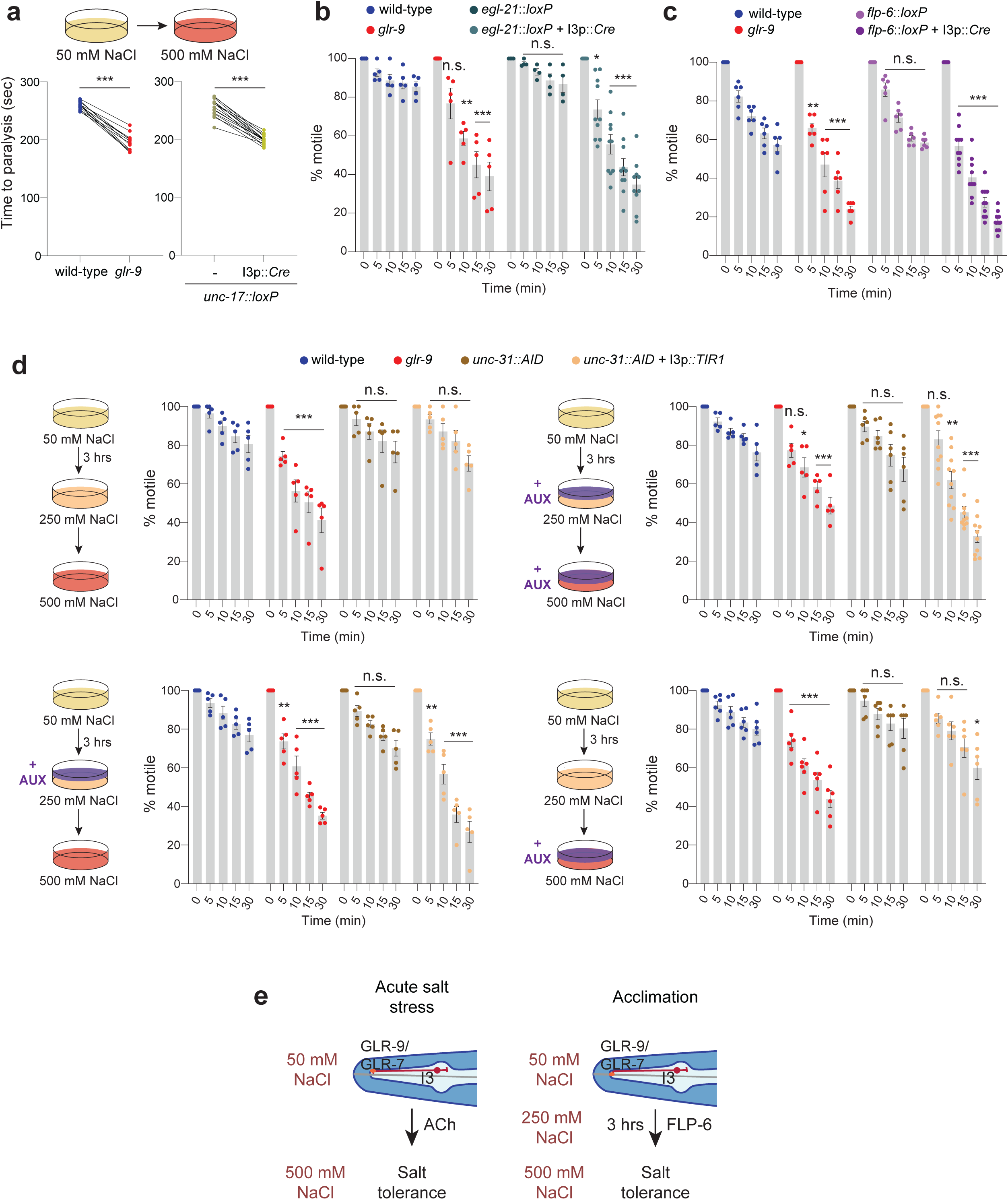
Cholinergic and FLP-6-mediated peptidergic signaling from I3 are necessary for high salt tolerance in unacclimated and acclimated animals, respectively. **a)** Quantification of time to paralysis upon shifting animals of the indicated genotypes from 50 to 500 mM NaCl (cartoon at top). Each dot is the time at which all ten animals in a single assay are immotile. Cre was expressed in I3 under the *glr-9* promoter. Control and mutant animals were examined in parallel in the same assay. *** indicates different from wild-type at P<0.001 (t-test). **b,c)** Percentage of animals of the indicated genotypes that are motile at each time point following a shift from the acclimation plate (250 mM NaCl for 3 hrs) to 500 mM NaCl. Each dot is the value from a single assay; n=30 animals per assay. Cre was expressed in I3 under the *glr-9* promoter. *, **, and *** indicate different from wild-type at the corresponding time at P<0.05, 0.01, and 0.001 (one-way ANOVA and Dunnett’s test). **d)** (Left in all panels) Cartoon of the experimental protocol. (Right in all panels) Percentage of animals of the indicated genotypes that are motile at each time point following a shift from the acclimation plate to the assay plate in the presence or absence of added exogenous auxin as shown. TIR1 was expressed in I3 under the *glr-9* promoter. Each dot is the value from a single assay; n=30 animals per assay. *, **, and *** indicate different from wild-type at the corresponding time at P<0.05, 0.01, and 0.001 (one-way ANOVA and Dunnett’s test). **e)** Cholinergic signaling from I3 promotes high salt tolerance under conditions of acute salt stress, whereas FLP-6 signaling from I3 during acclimation enables tolerance upon subsequent exposure to high salt. The *glr-9(oy180)* allele was used in all experiments. Horizontal lines in scatter plots indicate the mean; errors are SEM. Data shown are from 2-3 independent experiments each. n.s--not significant.

We next addressed the mechanisms by which GLR-9-mediated signaling contributes to salt acclimation. I3 responses to 500 mM NaCl were similar in animals with or without prior acclimation (Fig. S4a), indicating that acclimation does not alter I3 responses to high salt. Loss of *unc-17* specifically in I3 also did not affect acclimation (Fig. S4b), suggesting that cholinergic signaling from I3 regulates salt tolerance only under unacclimated conditions. Since I3 expresses multiple neuropeptides (Taylor et al., 2021), we next knocked out the *egl-21* carboxypeptidase required for processing of a subset of neuropeptides (Jacob and Kaplan, 2003) specifically in I3 using Cre-Lox mediated recombination (Huang et al., 2023) and assessed acclimation. Animals lacking *egl-21* in I3 became immotile more rapidly in high salt similar to *glr-9* mutants following acclimation (Fig. 4b), but responded similarly to wild-type animals upon acute salt challenge (Fig. S4c). We conclude that peptidergic signaling from I3 is required for acclimation and subsequent tolerance of high salt but is not required under conditions of acute salt stress.

Neuronal transcriptomics analyses indicate that I3 expresses at least 24 neuropeptide genes at different levels (Taylor et al., 2021). Analyses of salt acclimation phenotypes of a subset of I3-expressed neuropeptides showed that animals mutant for the *flp-6* FMRFamide-like neuropeptide failed to tolerate high salt following acclimation (Fig. S4d-e). The salt tolerance phenotype of *glr-9; flp-6* double mutants was similar to that of each single mutant alone suggesting that these molecules act in a linear pathway to regulate acclimation (Fig. S4e).

However, *flp-6* mutants behaved similarly to wild-type animals upon acute high salt exposure (Fig. S4f). Since *flp-6* is expressed in multiple neuron types in addition to I3 (Kim and Li, 2004; Taylor et al., 2021), we confirmed that *flp-6* acts in I3 by knocking out *flp-6* specifically in I3 via Cre-Lox-mediated recombination. Loss of *flp-6* expression specifically in I3 was again sufficient to reduce tolerance of high salt following acclimation similar to the phenotype of *flp-6* global mutants, but did not affect tolerance in an acute high salt challenge (Fig. 4c, Fig. S4f). FLP-6 has been shown to interact with multiple candidate neuropeptide receptors that are expressed broadly in neuronal and non-neuronal cells (Beets et al., 2023). However, mutations in single candidate receptor genes had minor or no effects on salt acclimation (Fig. S4g) suggesting that FLP-6 receptors may act redundantly within a single cell or across multiple cell/tissue types to regulate salt tolerance. Together, these results indicate that FLP-6-mediated signaling from I3 is necessary to promote tolerance of high salt following acclimation.

We next examined the temporal requirement for neuropeptide signaling from I3 to regulate salt tolerance following acclimation. To temporally modulate neuropeptide release from I3, we expressed the auxin receptor TIR1 specifically in I3 in a strain in which the *unc-31* CAPS-related protein required for dense core vesicle fusion (Speese et al., 2007) is endogenously tagged with degron sequences (Cornell et al., 2022; Nishimura et al., 2009; Zhang et al., 2015).

Addition of exogenous auxin to the growth plates during acclimation and exposure to high salt resulted in rapid paralysis further supporting the necessity of peptidergic signaling from I3 for tolerance of high salt (Fig. 4d). Addition of auxin only during the 3 hr acclimation period but not during high salt exposure was sufficient to induce rapid paralysis (Fig. 4d), indicating that peptidergic signaling from I3 is necessary only during the acclimation period to promote high salt tolerance. Taken together, our results indicate that cholinergic signaling from I3 increases tolerance of acute salt stress (Fig. 4e). In contrast, FLP-6 neuropeptide signaling from I3 during acclimation is protective in response to subsequent high salt exposure (Fig. 4e).

### Salt acclimation alters gene expression via both GLR-9-dependent and -independent pathways

The rapidity with which *C. elegans* loses fluid and paralyzes in hypertonic solutions is in part determined by the permeability of the animal’s cuticle as well as the concentration of internal osmolytes. Consistently, increased salt resistance following acclimation occurs as a consequence of altered expression of molecules such as cuticular collagens and osmolyte biosynthetic enzymes predicted to be protective against hypertonicity (Burkewitz et al., 2012; Dodd et al., 2018; Lamitina et al., 2004; Lamitina et al., 2006; Rohlfing et al., 2010). We hypothesized that I3 signaling regulates the expression of a subset of genes in distal tissues such as the cuticle, epidermis and gut that contribute specifically to salt tolerance, and that failure to correctly regulate these genes in *glr-9* mutants contributes to their decreased salt resistance. To test this notion, we compared the transcriptional profiles of wild-type and *glr-9* mutants prior to and following 3 hrs of salt acclimation (Fig. 5a-b).

**Fig. 5.**
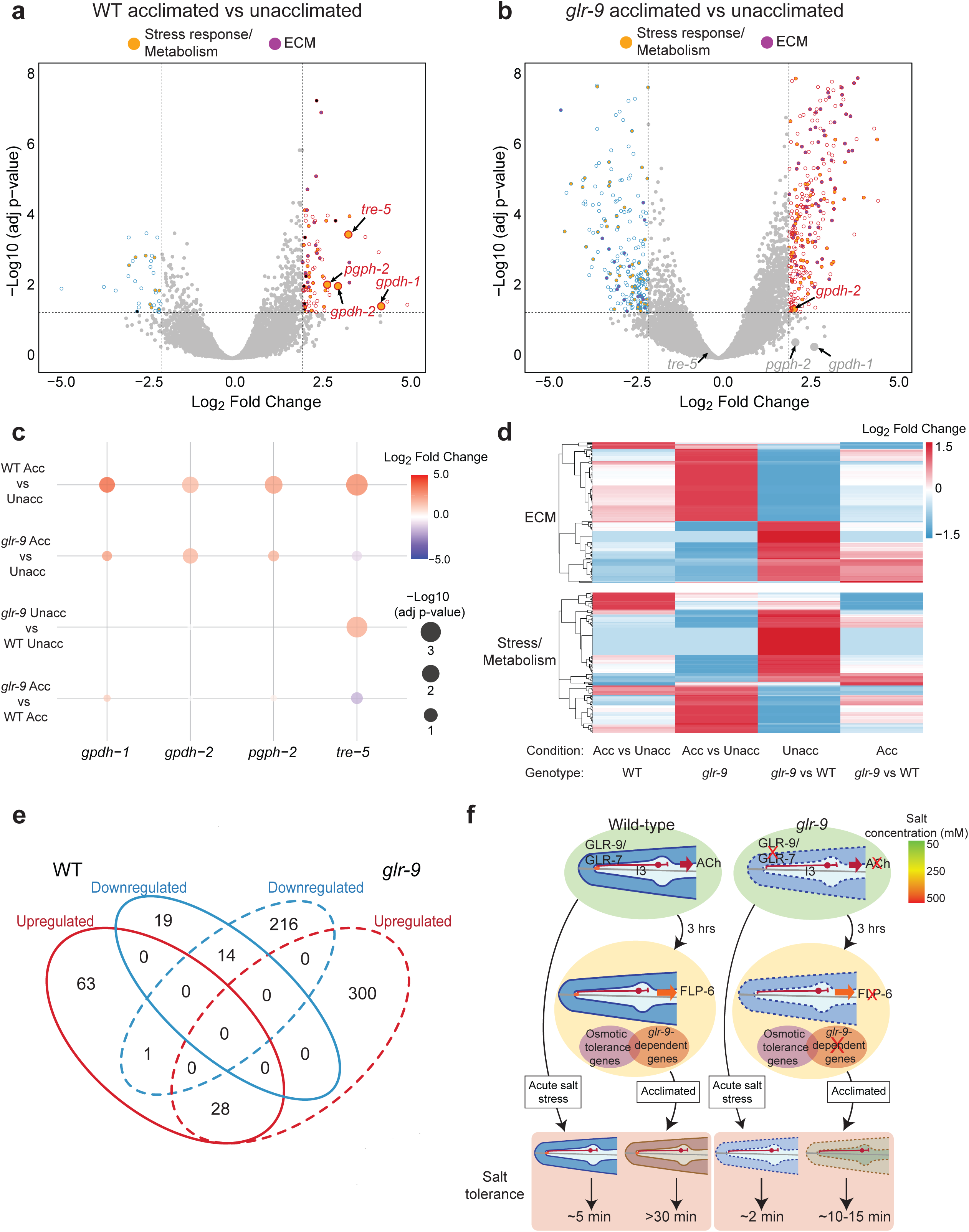
GLR-9-mediated signaling is required for altered expression of a subset of osmotolerance genes. **a,b)** Volcano plots of differential gene expression in unacclimated (50 mM NaCl) and salt acclimated (250 mM NaCl for 3 hrs) wild-type and *glr-9(oy180)* animals. Horizontal and vertical dashed lines indicate significant at adjusted p-value <0.05 and log_2_ fold change >2 or <−2, respectively. A subset of osmolyte biosynthetic enzymes previously implicated in osmotolerance is indicated. Differentially expressed genes predicted to encode molecules involved in stress responses or metabolism, and cuticular components, are indicated in yellow and purple, respectively. **c)** Dot plot of average expression changes of the indicated genes in the shown genetic backgrounds and conditions. Color indicates log_2_ fold change and size indicates adjusted p-value. Blank indicates no expression change detected. **d)** Hierarchically clustered heatmaps of differentially expressed genes categorized as being involved in extracellular matrix remodeling (top) or in stress and metabolic responses (bottom) in the indicated conditions and genetic backgrounds. **e)** Venn diagram indicating unique and common differentially expressed genes in unacclimated and acclimated wild-type and *glr-9(oy180)* mutants. **f)** Working model of the contribution of GLR-9-mediated signaling to high salt tolerance. In the absence of *glr-9* or cholinergic signaling from I3, remodeling of the cuticle even under isotonic conditions may contribute to decreased resistance to acute high salt stress. Upon acclimation, both GLR-9-dependent and -independent pathways contribute to cuticular remodeling and altered expression of osmotolerance genes, thereby enabling prolonged tolerance of high salt. The failure to alter expression of a subset of osmotolerance genes in *glr-9* mutants or upon loss of FLP-6 signaling from I3, and/or the dysregulated gene expression program under isotonic and acclimated conditions, reduces high salt tolerance following acclimation.

Principal component analyses indicated that transcriptional profiles clustered based on both genotype and condition (Fig. S5a). Comparison of the transcriptional profiles of wild-type animals prior to and following acclimation identified 92 and 33 significantly up- and down-regulated genes, respectively (Fig. 5a, Data Table 1). A small subset of these differentially expressed genes overlapped with those identified previously using partly distinct experimental conditions (Dodd et al., 2018; Rohlfing et al., 2011) (Fig. S5b). Consistent with previous observations, the *gpdh-1* and *gpdh-2* glycerol-3-phosphate dehydrogenase, *pgph-2* phosphoglycolate phosphatase, and *tre-5* trehalase encoding genes were upregulated following acclimation (Fig. 5a,c, Data Table 1). Transcriptional upregulation of these enzymes upon hypertonic stress contributes to the production of the osmolyte glycerol which is protective for hypertonic stress (Dodd et al., 2018; Lamitina et al., 2004; Lamitina and Strange, 2005; Possik et al., 2022; Rohlfing et al., 2011). The expression of a subset of genes encoding collagen was also altered consistent with a role for cuticular components in regulating hypertonic stress responses (Fig. 5a,d, Data Table 1) (Choe, 2013; Dodd et al., 2018; Lamitina et al., 2006; Rohlfing et al., 2011; Solomon et al., 2004; Urso and Lamitina, 2021; Wheeler and Thomas, 2006; Wimberly and Choe, 2022).

Acclimation resulted in altered expression of a larger set of genes in *glr-9* mutants (Fig. 5b, Data Table 1). However, the expression of *gpdh-1*, *pgph-2* and *tre-5* was no longer significantly altered in *glr-9* mutants (Fig. 5b-c, Data Table 1), indicating that I3-mediated signaling in response to salt is necessary for the upregulation of these enzymes. Although a subset of genes including genes encoding cuticular collagens was regulated similarly in both wild-type and *glr-9* mutants following acclimation (Fig. 5e, Data Table 1), the expression of distinct sets of genes implicated in stress responses and in the regulation of metabolism, as well as genes are predicted to encode components of the cuticle or hypodermis, was altered in acclimated *glr-9* mutants (Fig. 5b,d, Fig. S5c, Data Table 1). We conclude that altered gene expression upon salt acclimation is mediated via both I3-dependent pathways that may be salt-specific, and I3-independent pathways that may respond to general hypertonic stress.

We noted that the transcriptional profile of *glr-9* mutants was distinct from those of wild-type animals even at 50 mM NaCl (Fig. 5c-d, Fig. S5a,d, Data Table1) suggesting that chronic I3-mediated signaling may be necessary for regulating physiology even at isotonic salt concentrations. Significantly upregulated genes in *glr-9* mutants included genes implicated in the regulation of metabolism and stress, multiple components of the cuticle and hypodermis including collagen and chitinase-like genes, as well as nematode-specific groundhog-like (GRL) and ground-domain (GRD) secreted proteins that are apical extracellular matrix components of the cuticle and precuticle (Hao et al., 2006; Serra et al., 2024) (Fig. 5b,d, Fig. S5d, Data Table 1). These observations suggest that chronic *glr-9* signaling under standard growth conditions may signal the presence of salt, and that the cuticular barrier may be structurally remodeled in the absence of this signaling, potentially contributing to the decreased resistance of these animals to acute salt stress (see Discussion).

To assess the functional significance of these expression changes, we tested two predictions arising from the transcriptional profiling data. Animals moved from 50 mM NaCl to either 400 mM or 500 mM NaCl exhibit similar initial responses including rapid shrinkage and paralysis (Choe, 2013; Choe and Strange, 2007a). However, while prolonged exposure to 500 mM NaCl leads to death (Lamitina et al., 2004; Solomon et al., 2004), animals regain body volume and motility after 2-4 hrs on 400 mM NaCl via modulation of ion transport and upregulation of osmotolerance genes (Choe and Strange, 2007a, b). The failure of *glr-9* mutants to upregulate osmotolerance genes and their extensively remodeled cuticle even under isotonic conditions predicts that these animals will be unable to recover, or will recover more slowly, on 400 mM NaCl. Consistent with this prediction, we found that *glr-9* mutants regained motility significantly more slowly than wild-type animals under these conditions (Fig. S6a). These results further support the notion that I3 signaling-regulated gene expression changes play a critical role in regulating salt tolerance.

The finding that salt acclimation results in gene expression changes that are partly I3-dependent and are thus salt-specific, also predicts that different hypertonic solutions may alter the expression of different gene subsets, with only a shared subset being regulated by general osmotic stress. This hypothesis predicts that acclimation to high salt would not be protective in response to high sugar challenge and *vice versa*. We found that acclimation to 250 mM NaCl did not increase tolerance of either wild-type or *glr-9* mutants to 500 mM sucrose, and conversely, acclimation to 250 mM sucrose had no effect on resistance to 500 mM NaCl under our experimental conditions (Fig. S6b). We infer that partly distinct pathways contribute to the detection of individual hypertonic solutes to regulate specific organismal stress responses.

## DISCUSSION

Here we show that salt-sensing by the single I3 enteric neuron specifically promotes salt stress resistance in *C. elegans.* I3 salt responses are dependent on the GLR-9/GLR-7 IR proteins which are both necessary and sufficient for salt detection. I3 signals via cholinergic and peptidergic signaling to increase salt stress resistance in unacclimated and salt-acclimated animals, respectively. We find that signaling from I3 modulates the expression of genes that regulate cuticle composition and internal osmolyte concentration, resulting in tolerance of high salts but not high sugars (summarized in Fig. 5f). Our observations describe the molecular and neuronal mechanisms by which chemosensation by an enteric neuron regulates physiological homeostasis in response to a specific abiotic stress.

Chemoreceptors are among the fastest evolving gene families in multicellular animals, with some gene families being entirely lost or gained within specific lineages (Policarpo et al., 2024; Sanchez-Gracia et al., 2009; Valencia-Montoya et al., 2024). For example, while variant IR sensory proteins are found in all protostomes, they are absent from deuterostomes (Croset et al., 2010; Valencia-Montoya et al., 2024). However, to date, chemosensory roles for these IR proteins have been described and/or suggested primarily in arthropods (Benton et al., 2009; Eyun et al., 2017; Graeve et al., 2022; Groh-Lunow et al., 2014; Ni, 2020). Our identification of a chemosensory function of an IR in *C. elegans* implies that IRs may have retained chemosensory functions in the protostome lineage through evolution. It remains to be determined whether the remaining PEN-expressed IRs in *C. elegans* also function as chemosensors, or whether similar to some *Drosophila* and mosquito IRs (Ni, 2020; van Giesen and Garrity, 2017), subsets of these proteins also detect other sensory cues.

GLR-9-mediated salt sensation is transmitted via distinct signaling pathways from I3 to modulate gene expression in tissues and organs such as the gut and cuticle to enhance salt tolerance. We find that even under isotonic conditions, *glr-9* mutants exhibit broad changes in the expression of genes that may impact hypertonic salt resistance. Since I3 responds tonically to a range of salt concentrations, we propose that chronic activation of GLR-9 and cholinergic signaling from I3 upon growth on low salt enhances resistance to acute high salt. Remodeling of the cuticle in the absence of I3 signaling may thus sensitize animals to high salt stress, resulting in more rapid paralysis upon an acute high salt challenge (Fig. 5f). During acclimation, FLP-6-mediated signaling from I3, as well as from other neuronal and non-neuronal osmosensory pathways, drive broad gene expression changes that promote salt tolerance. The failure to alter the expression of a subset of these genes, including osmolyte biosynthesis genes, in acclimated *glr-9* mutants, along with their dysregulated gene expression under isotonic conditions, may underlie their decreased salt tolerance phenotype (Fig. 5f). Additional pathways likely also contribute to osmotolerance under different conditions and timepoints following stress exposure (Huang et al., 2007; Igual Gil et al., 2017; Lee et al., 2016).

While exposure to hypertonic solutions can cause general effects such as altered protein quality control and tissue deformation (Burkewitz et al., 2011; Choe and Strange, 2008; Lamitina et al., 2006; Urso and Lamitina, 2021), several lines of evidence including our findings suggest that *C. elegans* and other animals can discriminate among different hypertonic solutes. First, equiosmolar concentrations of salt and sucrose result in distinct survival rates in *C. elegans* (Chandler-Brown et al., 2015; Choe and Strange, 2007a; Lamitina et al., 2004), implying that individual osmotic stressors can elicit partly different organismal responses. Second, *glr-9* mutants exhibit decreased tolerance of high salts but not equiosmolar sugar, highlighting the salt-specificity of the I3 pathway. Third, acclimation to NaCl does not confer resistance to sucrose and *vice versa* under our experimental conditions (see (Chandler-Brown et al., 2015)), indicating that the adaptive response to one osmotic stressor is not fully sufficient to protect against others. Fourth, hypertonic salt regulates the expression of a subset of osmotolerance genes in a GLR-9-dependent manner, further suggesting the presence of solute-specific pathways. Similarly, in rodents, while a number of physiological responses are independent of hyperosmotic solute identity, arterial blood pressure regulation and sympathetic nerve activity are more sensitive to central NaCl infusion than to equiosmolar sugars (Kinsman et al., 2017). Moreover, subsets of OVLT neurons in the circumventricular organs of the brain preferentially respond to hypertonic NaCl over equiosmolar sugars (Kinsman et al., 2017; Stocker et al., 2024), indicating the existence of NaCl-specific, non-osmosensory pathways. Given the critical role of salts in regulating physiological processes, we suggest that dedicated neuronal pathways for high salt detection provides additional flexibility in coordinating responses based on internal state.

Enteric sensory neurons in *C. elegans* appear to utilize partly distinct sensory mechanisms from those of somatic exteroceptive sensory neurons. For instance, the GUR-3 gustatory receptor-like protein required for hydrogen peroxide detection by the I2 PEN in *C. elegans*, and the DEL-3 and DEL-7 ASIC channels that detect ingested bacteria in the NSM PEN are expressed exclusively or primarily in PENs (Bhatla and Horvitz, 2015; Rhoades et al., 2019; Taylor et al., 2021). Similarly, 6 of the 7 IRs including GLR-7 and GLR-9 are expressed primarily or highly in PENs. Moreover, PENs are non-ciliated unlike exteroceptive sensory neurons, suggesting novel mechanisms for receptor trafficking and localization at their sensory endings. The intrinsic enteric neuronal network has ancient evolutionary origins and is thought to have arisen prior to and independently of the central nervous system (Furness and Stebbing, 2018; Watanabe et al., 2009). Investigating the mechanisms underlying enteric neuron development and function in diverse organisms may provide insights into ancestral neuronal strategies, and expand our understanding of interoceptive sensory functions across species.

## METHODS

### Growth of *C. elegans* strains

*C. elegans* strains were grown on NGM plates seeded with *E. coli* OP50 at 20°C. *unc-122*p*::dsRed* or *gfp* were used as co-injection markers (at 40 ng/μl) to generate transgenic strains. Strains were generated using standard genetic crosses. The genotypes of all strains were verified by sequencing or phenotypic examination. All experiments were performed using young adult hermaphrodites grown under well-fed conditions for at least three generations. A list of all strains used in this work is provided in Table S1.

### Molecular biology

1.1 kb of sequences upstream of and including the *glr-9* START codon were amplified from wild-type animal lysate and used to drive expression specifically in I3. Expression in I1 was driven using 1.1 kb of sequences upstream of the *lgc-8* START codon. GLR-9 cDNA was amplified from wild-type animal lysate and tagged with *gfp* for localization and for functional rescue experiments. Since *glr-7* has *a* and *b* isoforms, the genomic region of *glr-7* from the START codon of the *b* isoform (X: 2,419,477) to the STOP codon of both *a* and *b* isoforms (X:2,414,064) was amplified from wild-type animal lysate. These sequences were tagged with *TagRFP* to examine localization of *glr-7*. All plasmids were generated using standard cloning methods and verified by Sanger sequencing. A list of plasmids used in this work is provided in Table S2.

### CRISPR/Cas9-mediated genome editing

Deletion alleles of *glr-9* and *osm-9* were generated using an injection mix containing 0.5 μl of Cas9 (IDT), 1.5 μl of 100 μM tracrRNA, 1 μl of 50 μM 5’-crRNA, 1 μl of 50 μM 3’-crRNA, 1 μl of 17 μM *dpy-10* crRNA as a co-CRISPR injection marker, and nuclease-free water, for a total volume of 10 μl. No DNA donor was used. Worms were maintained at 25°C after injection. Three days following injection, dumpy or roller F1 progeny were segregated onto individual plates. After the singled F1 animals laid eggs, parents were lysed and genotyped using PCR.

To generate the *flp-6::loxP* insertion strain (PY12288, Table S1), the injection mix included 0.5 μl of Cas9 (IDT), 2.8 μl of 34 μM crRNA, 0.2 μl of 100 μM tracrRNA, 2 μl of 26 μM donor DNA, 50 ng/μl of column-purified *unc-122*p*::gfp*, and nuclease-free water for a total volume of 20 μl. Following injection, worms were kept at 25°C. F1 progeny carrying the *unc-122*p*::gfp* co-injection marker were singled onto individual plates and genotyped. The C-terminal *loxP* site was inserted first, followed by insertion of the N-terminal *loxP* site. All tracrRNA, crRNAs, and donor DNAs were obtained from IDT. The sequences of the crRNAs and donor DNA are listed below.

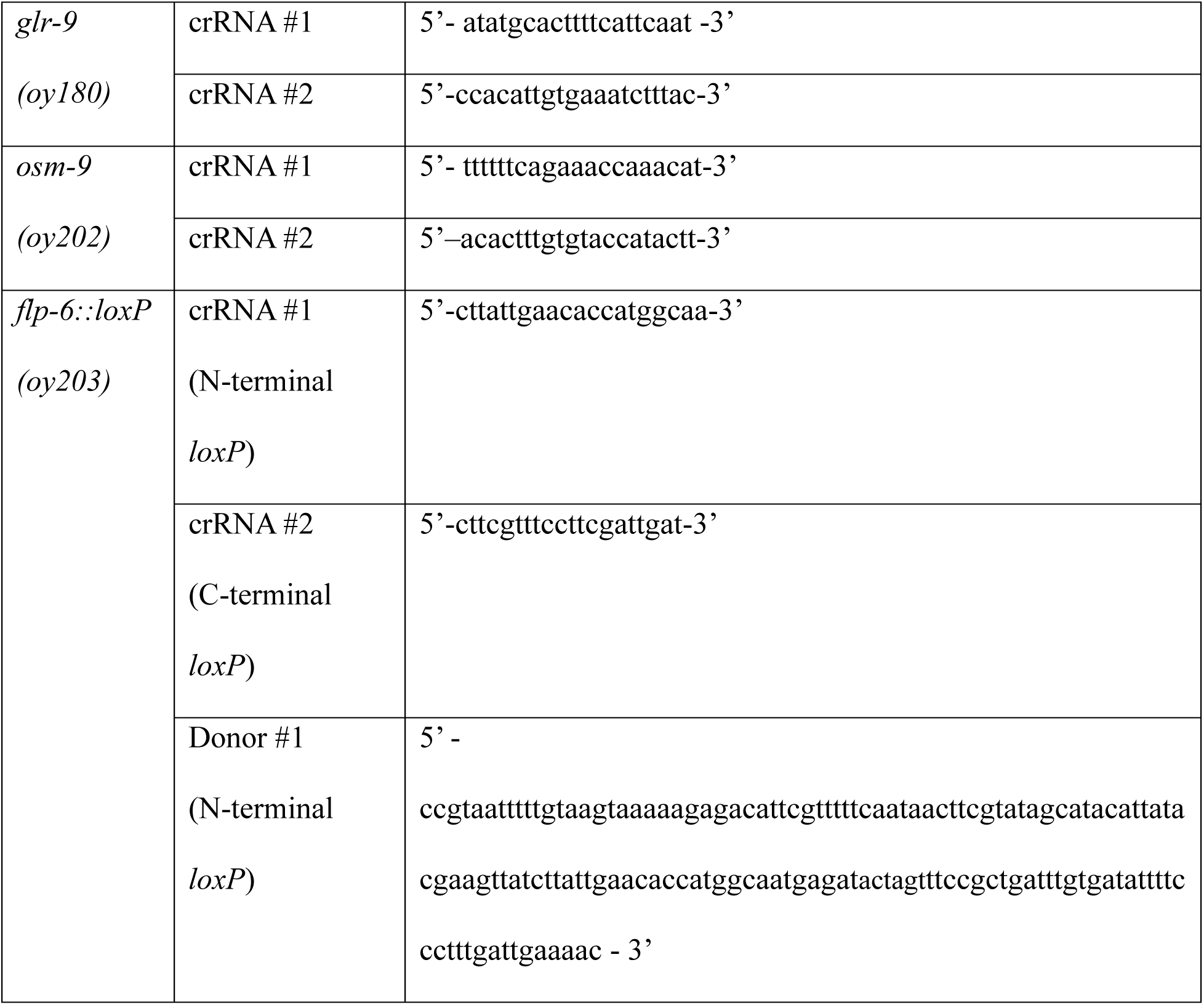
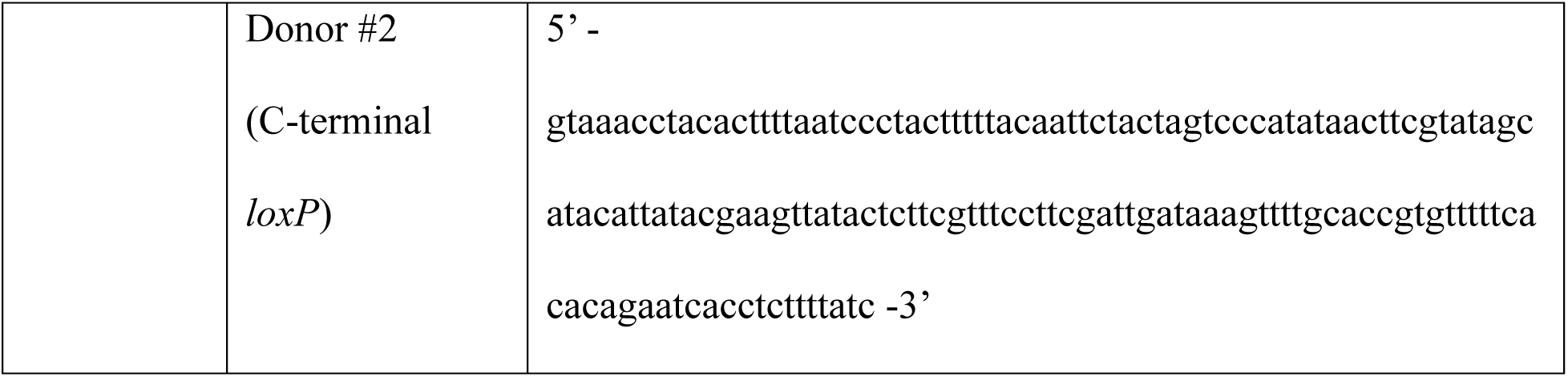

### Phylogenetic analysis

iGluR sequences from *Mus musculus* and *Drosophila melanogaster* were obtained from Croset et al (Croset et al., 2010). iGluR and IR sequences from *C. elegans* were obtained from Wormbase (www.wormbase.org). Sequences were aligned using MUSCLE (Edgar, 2004), and a phylogenetic tree was constructed using the Maximum Likelihood method with adaptive bootstrapping (116 replicates) in MEGA 12.0.9. The resulting tree was visualized and modified using FigTree v1.4.4.

### Fluorescent reporter imaging

1 day-old adult animals were anesthetized with 10 mM (-)- tetramisole hydrochloride (Sigma-Aldrich L9756) and mounted on 10% agarose pads on microscope slides. Images in Figure 1b,d, Fig. S2i were acquired at 0.2 μm *z*-intervals using a 100× oil immersion objective on an inverted two-camera spinning disk confocal microscope (Leica DMI6000B with a CSU-W1 spinning disk head and two Andor Neo sCMOS cameras) and Andor IQ3.5 Software. Images in Figure 1e and Fig. S2j were acquired using a 64× oil immersion objective on an inverted spinning disk confocal microscope (Zeiss AxioObserver with Yokogawa CSU-X1 confocal head system and QuantEM SC512 camera) and SlideBook^TM^ software. All images were processed using FIJI/Image J.

### 3D reconstruction of I3 sensory ending

Serial section electron microscope (EM) images of I3 were obtained from Wormimage (www.wormimage.org). I3 was identified as “blue 14” on the EM images (with the assistance of David Hall), specifically from the sections N2T_114522 to N2T_114583. The TrakEM2 plugin in Fiji was used for manually aligning I3 images. Following the alignment, manual segmentation was performed using 3dmod v4.11 from the IMOD suite (https://bio3d.colorado.edu/imod/) to generate the 3D reconstruction of the I3 sensory ending.

### Acute salt stress assay

Acute responses to high salt were examined using a protocol adapted from (Rohlfing et al., 2010). 6 cm unseeded NGM plates supplemented with the indicated salt concentrations were divided into two sections to assess the behaviors of wild-type and *glr-9* mutant in parallel. Growth-synchronized 1-day old adult hermaphrodites were transferred from NGM plates containing 50 mM NaCl to high salt-containing plates to initiate the assay. Each assay included 10 animals of each genotype, and the time at which all 10 animals were unresponsive to touch with a platinum wire was recorded. Unresponsive was defined as failure to move upon prodding with a platinum wire at the nose and tail, and cessation of pharyngeal pumping. At least three independent assays were conducted over multiple days for each data point.

### Salt acclimation assay

Acclimation experiments were performed using an adaptation of a previously published protocol (Rohlfing et al., 2010). 35 growth-synchronized 1-day adult hermaphrodites were transferred from NGM growth plates (50 mM NaCl) to NGM plates containing 250 mM NaCl, 250 mM KPO_4_, or 250 mM sucrose and seeded with 200 μl *E. coli* OP50. Animals were maintained at 20°C for 3 hrs. Following 3 hrs of acclimation, animals were transferred to unseeded plates containing 500 mM NaCl, 500 mM KPO_4_, or 500 mM or 1 M sucrose, and tested for responses upon prodding with a platinum wire after 5, 10, 15 and 30 mins. Unresponsive was defined as failure to move upon prodding with a platinum wire at the nose and tail, and cessation of pharyngeal pumping. At least three independent assays were conducted over multiple days for each data point.

### Salt recovery assay

Recovery from paralysis on 400 mM NaCl was examined using an adaptation of previously published protocols (Choe and Strange, 2007a; Lamitina et al., 2004). Bacteria-seeded growth plates were supplemented with 400 mM NaCl and divided into two sections to assess wild-type and *glr-9* animals in parallel. Growth-synchronized 1-day old adult hermaphrodites were transferred from growth plates containing 50 mM NaCl to the assay plates and were examined every 15 mins up to 4 hrs. Each assay included 10 animals of each genotype. Unresponsive was defined as failure to move upon prodding with a platinum wire at the nose and tail, and cessation of pharyngeal pumping. At least three independent assays were conducted over multiple days for each data point.

### Glycerol avoidance assay

The avoidance assay was performed as previously described (Culotti and Russell, 1978). Briefly, a ring of 8M glycerol with xylene cyanol (Sigma-Aldrich X4126) was pipetted onto an unseeded NGM plate dried overnight. For each assay, 10 one-day-old adult hermaphrodites were transferred to the center of the glycerol ring. After 10 mins, the number of animals inside and outside the ring was counted. Wild-type and *glr-9*(*oy180)* mutants were examined in parallel.

### Auxin-induced degradation

400 mM Auxin [1-naphthaleneacetic acid (NAA), Sigma 317918)] was dissolved in EtOH and added to NGM containing the indicated salt concentrations to make 4 mM auxin-containing plates. Control plates contained equal concentrations of EtOH alone. Plates were seeded with 100 μl *E. coli* OP50, dried overnight, and stored at 4°C wrapped in aluminum foil to prevent light exposure.

At least 70 young adult hermaphrodites were transferred from NGM growth plates to plates containing 250 mM NaCl supplemented with either auxin or ethanol. After 3 hrs acclimation, animals were tested for movement on plates containing 500 mM NaCl supplemented with either auxin or ethanol. Animals were examined at 5, 10, 15, and 30 mins for responses to touch with a platinum wire. Each assay included 30 animals per condition and per genotype and was conducted over multiple days. At least 2 sets of experiments were performed for each genotype on the same day. Experimental and control strains were examined together in the same assays in each experiment.

### Quantification of pumping frequency

Pumping frequency was measured manually using 1-day old adult animals on OP50-seeded NGM plates containing 50 mM or 250 mM NaCl. For each data point, the pumping frequency of individual animals was measured three times for 20 sec each to obtain the average pumping frequency. Wild-type and *glr-9* mutants were examined together in the same assays for each experiment. At least three independent assays were conducted over multiple days for each data point.

### Microfluidics behavioral assay

Behavioral assays using microfluidics devices were performed essentially as previously described (Khan et al., 2022). After degassing the assembled microfluidic devices, the outlet port was loaded with 5% v/v poloxamer surfactant (Sigma P5556) containing 2 mg/ml xylene cyanol to eliminate bubbles. 20 to 30 1-day old adult hermaphrodites were transferred to unseeded NGM plates, washed with water, and loaded into two separate arenas within a single device. *glr-9* mutants were assessed in the same device, with wild-type controls placed in the adjacent arena. After allowing the worms to disperse (<5 min), the stimulus flow containing either 75 mM NaCl or 250 mM NaCl and 2 mg/ml xylene cyanol for visualization was initiated. 20 min videos were recorded at 2 Hz using a PixelLink camera. Microfluidic devices were cleaned after each experiment by washing with water and 100% ethanol before reuse. Custom MATLAB software was used to process and analyze all recordings, and data were visualized using custom R scripts (https://doi.org/10.5281/zenodo.13748735). Mean residency histograms and chemotaxis indices were analyzed as previously described (Khan et al., 2022). Three independent assays were performed over multiple days.

### Calcium imaging

Custom microfluidic devices for calcium imaging were fabricated as described previously (Khan et al., 2022). Images were acquired using an Olympus BX52WI microscope with a 40× oil objective and a Hamamatsu Orca CCD camera at 4 Hz with 4 × 4 binning. All stimuli were diluted in filtered Milli-Q water, and animals were paralyzed with 10 mM (-)-tetramisole hydrochloride (Sigma L9756) dissolved in water prior to loading on the unseeded NGM. Stimuli dilutions were prepared fresh in water on the day of each imaging session. Responses to 30 sec of water, 30 sec of stimuli, and 30 sec of water were recorded in I3. Data were collected from biologically independent experiments conducted over at least two days. For GCaMP fluorescence analysis, recorded videos were processed using Fiji as previously described (Khan et al., 2022). Briefly, images were aligned using the Template Matching plugin, and cell body and background fluorescence were calculated using manually drawn ROIs. Background-subtracted fluorescence intensity values were used for analysis in R Studio. The baseline fluorescence intensity (F_0_) was calculated as the average ΔF/F_0_ value over 5 sec prior to stimulus onset. R Studio was used to compute the response mean and SEM (https://doi.org/10.5281/zenodo.13748735). Peak ΔF/F_0_ values were calculated as the maximum change in fluorescence relative to F_0_ within 10 sec following stimulus addition.

### RNA-Seq

Animals were synchronized by bleaching adults, collecting eggs, and then arresting hatched L1s in the absence of food for ∼16 hours. ∼1,000 animals were cultured on *E. coli* OP50 seeded at 1 ml per 10 cm plate at 20°C until the 1 day-old adult stage. Animals from a total of four 10 cm plates were used per sample. Animals were washed with M9 containing 0.01% Triton (Sigma T8787) and placed onto either 10 cm OP50 seeded NGM plates or NGM plates supplemented with 250 mM NaCl for 3 hrs at 20°C. Animals were then washed off the plates with M9 containing 0.01% Triton. Total RNA extraction was performed using TRIzol (Invitrogen 15596026) and chloroform (Sigma 288306), followed by RNA Clean & Concentrator™-5 (Zymo Research R1014) to extract total RNA. The quality and concentration of total RNA was assessed using Agilent High Sensitive RNA ScreenTape analysis (Agilent 5067-5579).

100 ng/µl of total RNA per sample was used as input to generate a sequencing library using the xGen™ Broad Range RNA library preparation Kit (IDT 10009865) with AMPure XP Beads (Beckman Coulter A63880). Agilent DNA ScreenTape (D1000 5067-5582) assays were used to assess the quality of the library prior to sequencing. Libraries were paired-end sequenced at 75×75 bases on a NextSeq 1000 system (Illumina). Sequencing reads were adapter trimmed using Trim Galore (https://doi.org/10.5281/zenodo.7598955), mapped to the WBcel235 *C. elegans* genome, and counted using STAR (Dobin et al., 2013) with alignMatesGapMax 2500.

Differential expression analysis was performed in R studio with DESEQ2 (Love et al., 2014) (BioProject accession number PRJNA1227491). The PCA, heatmaps, and volcano plots were generated using the ggplot package in R. Venn diagrams were created using VennDiagram in R. GO terms, Wormbase, WomCat, Uniprot, and PANTHER were used to categorize individual genes listed in Data Table 1.

### Statistical analyses

Statistical analyses were performed using Prism v10.4.1. Comparisons to wild-type were performed with t-test or one-way ANOVA across the genotype and condition unless indicated otherwise. Corrections for multiple comparisons were performed using Dunnett’s tests. Scatter plots and bar graphs were generated and analyzed using Prism 10. The statistical tests used, significance values, and the number of animals and assays are indicated in the legend to each Figure.

## Data availability

Underlying data for all behavioral and calcium imaging experiments have been deposited to Figshare. All other data necessary for confirming the conclusions of this article are represented fully within the article and its tables and figures.

## Acknowledgements

We are grateful to David Hall for assistance with identification of I3 in electron microscopy serial sections, Kirsten Judge for assistance with imaging, and Evelyn Keefer, Gillian Berglund, and Katherine Abruzzi for help with RNA-Seq experiments. We thank Todd Lamitina, Steven Flavell, Ugur Dag, and Yung-Chi Huang for discussion and advice, and Paul Garrity, Steven Flavell, Nathan Harris, Todd Lamitina, Stephen Nurrish, and Alison Philbrook for critical comments on the manuscript. A subset of strains was obtained from the *Caenorhabditis* Genetics Center and the National BioResource Project (Japan). This work was funded in part by the NIH (R35 GM122463 – P.S., T32 MH 19929 – L.C., T32 GM139798 and F31 NS134251 – S.B.) and the NSF (IOS 2042100 – P.S.).

---

**Fig. S1.**
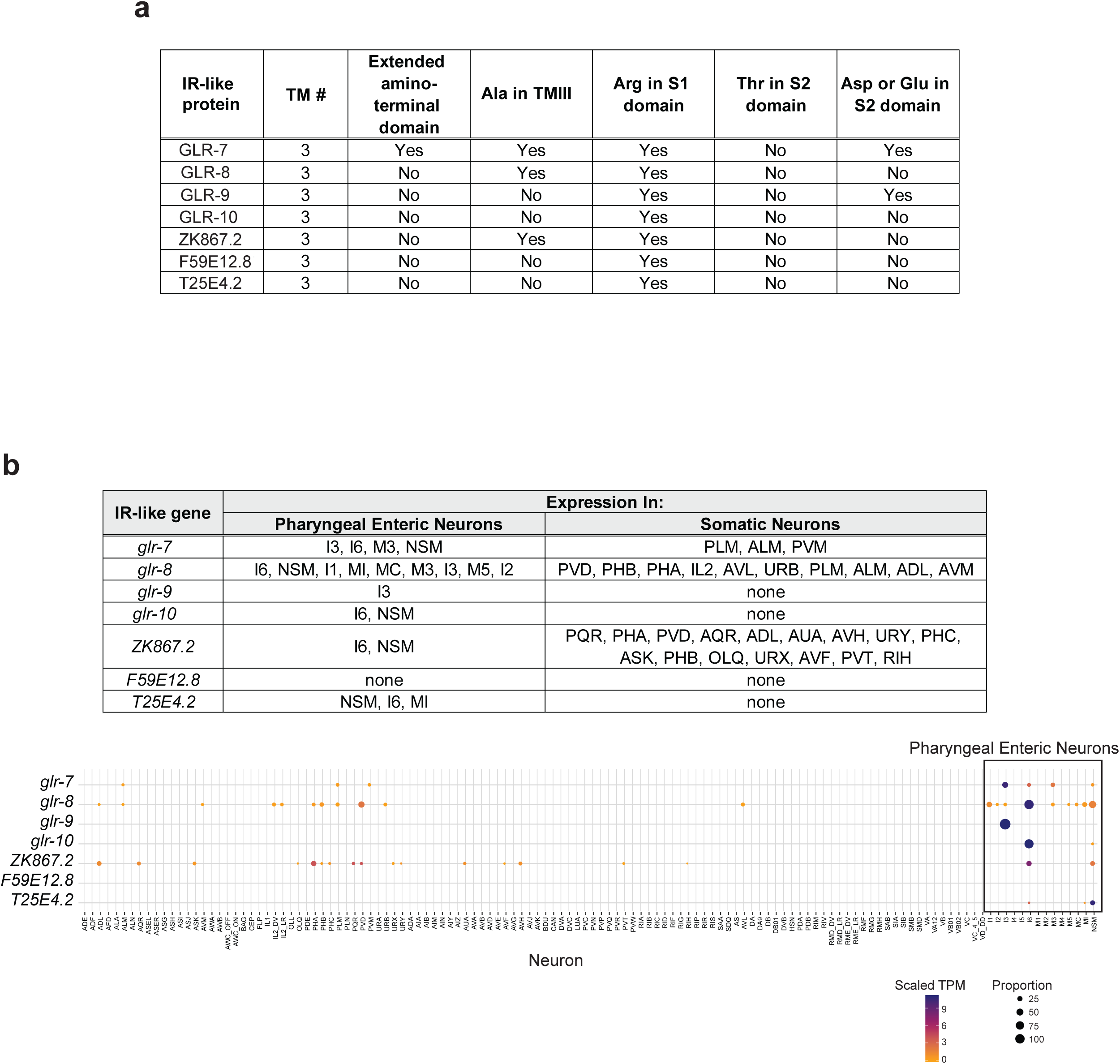
*C. elegans* IR-like genes are expressed highly or exclusively in PENs. **a)** Features of IR-like proteins in *C. elegans*. TM: transmembrane domain. Mutating the conserved Ala in TMIII in ionotropic glutamate receptors results in a constitutively open channel (Zheng et al., 1999; Zuo et al., 1997). The Arg, and Thr and Asp/Glu residues, in the S1 and S2 ligand-binding domains, respectively, directly interact with glutamate or synthesized agonists in ionotropic glutamate receptors (Benton et al., 2009). **b)** Predicted expression of IR-like genes adapted from the *C. elegans* neuronal gene expression map and network (CeNGEN) (Taylor et al., 2021).

**Fig. S2.**
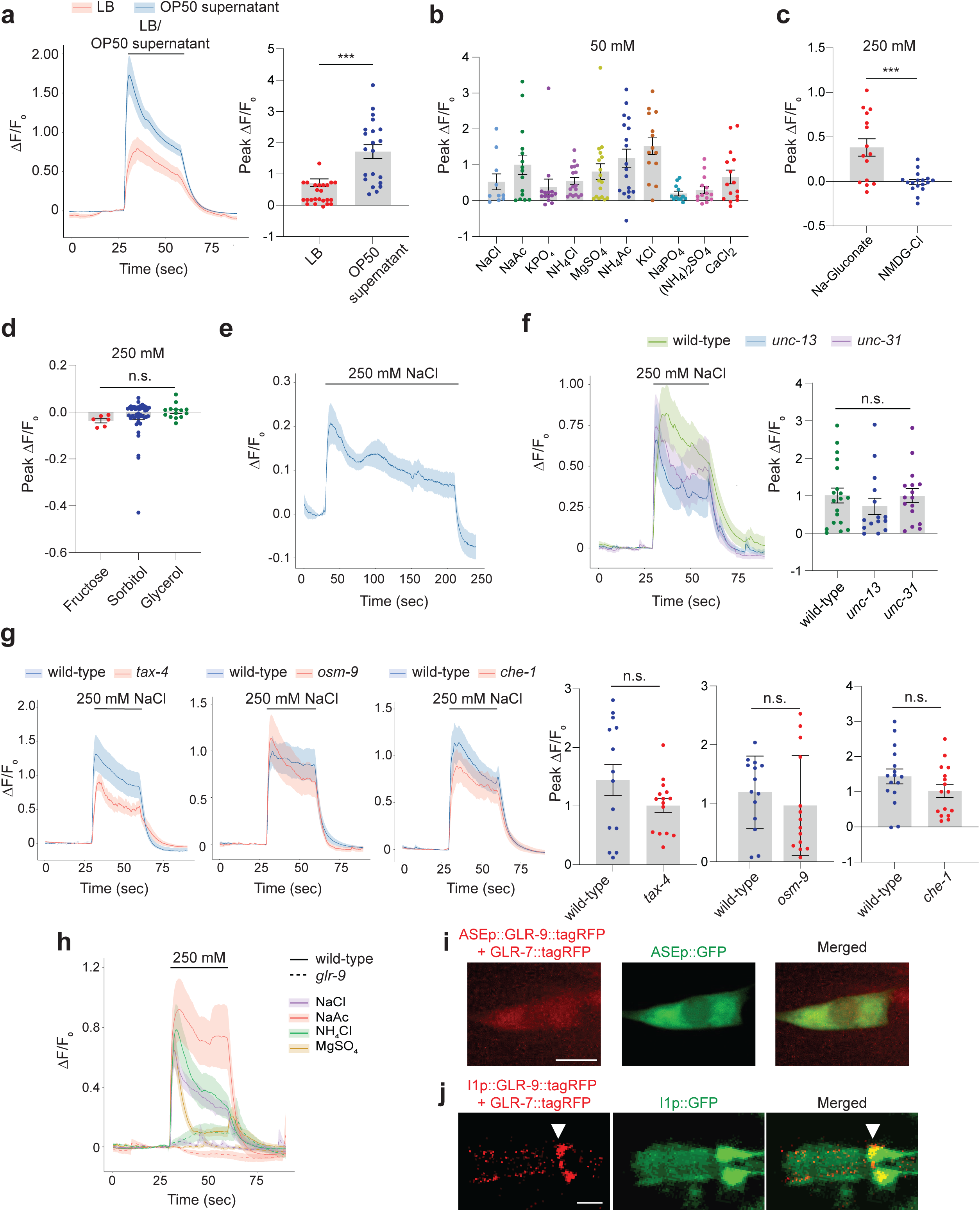
I3 responds cell-autonomously to multiple cations via GLR-9. **a)** Average (left) and peak intensity changes (right) of GCaMP6s fluorescence in wild-type I3 neurons in response to a 30 sec pulse of LB alone or supernatant of an OP50 bacterial culture grown in LB. *** indicates different at P<0.001 (t-test). **b-d)** Peak fluorescence intensity changes in I3 in response to the indicated chemical concentrations. Average traces are shown in Fig. 2a-c. *** indicates different at P<0.001 (c: t-test; d: one-way ANOVA and Dunnett’s test). **e)** Mean GCaMP6s fluorescence change in I3 in response to a 3 min pulse of 250 mM NaCl. **f,g)** Average (left) and peak intensity changes (right) of GCaMP6s fluorescence in I3 in response to a 30 sec pulse of 250 mM NaCl in animals of the indicated genotypes. Alleles used were *unc-13(e51), unc-31(e169)*, *tax-4(p678), osm-9(oy202),* and *che-1(p674)*. **h)** Mean GCaMP6s fluorescence changes in response to a 30 sec pulse of the indicated salts at 250 mM in wild-type and *glr-9(oy180)* mutants. n>17 each. Wild-type data were interleaved with data in Fig. 2d-g, and are repeated. **i,j)** Representative images of GLR-9/GLR-7::tagRFP localization in ASE soma (i), and I1 (j). Expression in ASE and I1 was driven under the *che-1* and *lgc-8* promoters, respectively. Arrowhead indicates the non-ciliated sensory ending of I1 in j. Anterior is at left in all images. Scale bar: 5 μm. Shaded regions in all traces indicate SEM. Each dot in the scatter plots is the value from a single neuron. Horizontal lines in scatter plots indicate the mean; errors are SEM. Data shown are from 2-3 independent experiments each. n.s.-not significant.

**Fig. S3.**
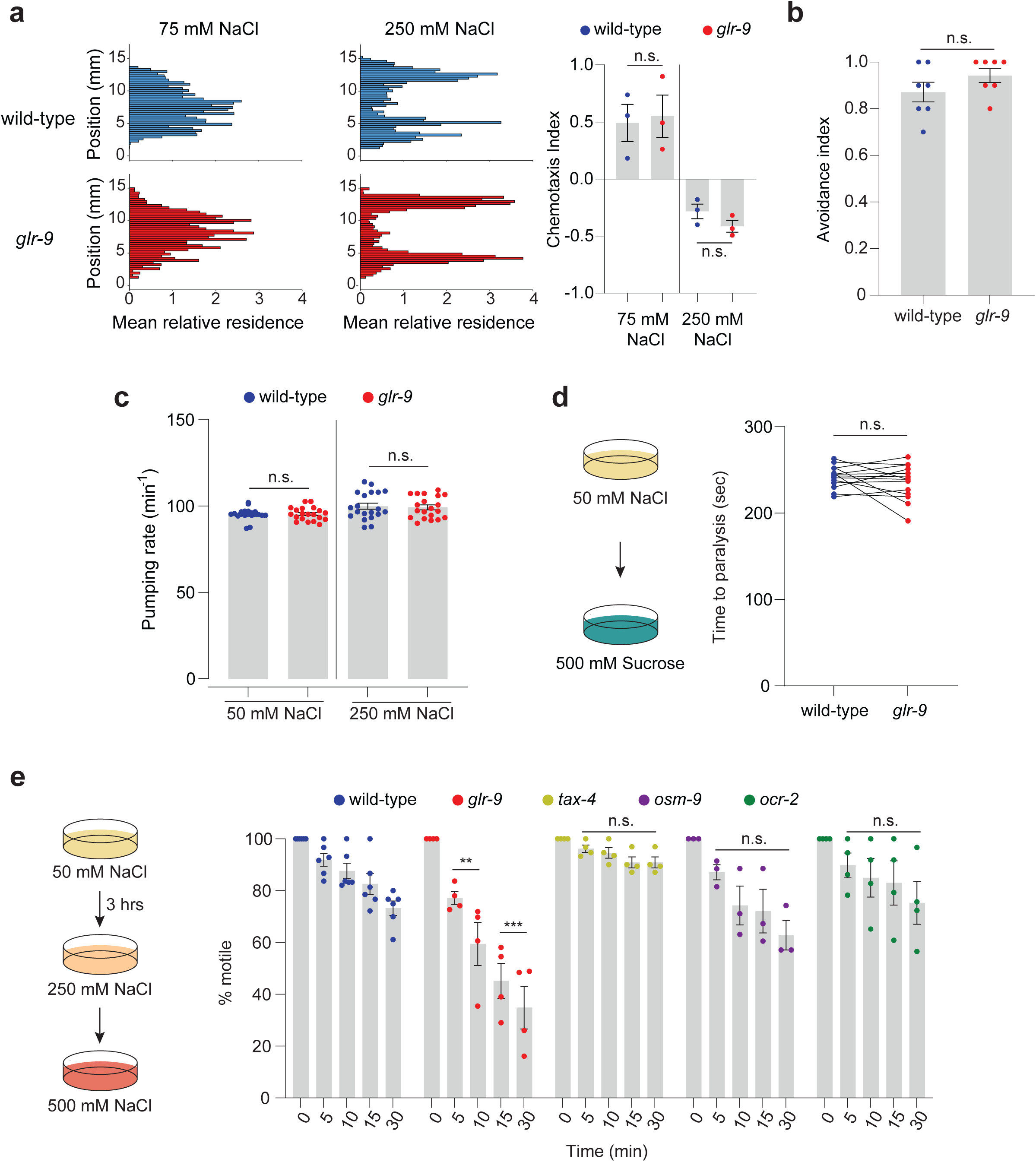
*glr-9* mutants do not exhibit altered behavioral responses to salt. **a)** (Left) Histograms showing average relative residence of wild-type and *glr-9(oy180)* animals in a microfluidics behavioral arena (Albrecht and Bargmann, 2011; Khan et al., 2022) with a central stripe of the indicated salt concentrations. (Right) Chemotaxis indices quantified from the microfluidics behavioral assays. Each dot is the index from a single assay of ∼15 animals each. **b)** Percentage of animals of the indicated genotypes remaining within the glycerol ring after 10 mins. Each dot is the value from a single assay of 10 animals each. **c)** Pumping rate on bacteria-seeded plates containing the indicated salt concentrations. Each dot is the measurement from a single animal. **d)** Quantification of time to paralysis of wild-type or *glr-9* mutants in the shown assay conditions (cartoons at left). Each dot is the time at which all ten animals in a single assay are immotile. Wild-type and *glr-9* mutants were examined in parallel in the same assay. **e)** Percentage of animals of the indicated genotypes that are motile at each time point following a shift from the acclimation plate to the assay plate. Alleles used were *glr-9(oy180), tax-4(p678), osm-9(ky10)*, and *ocr-2(ak47)*. Each dot is the value from a single assay; n=30 animals per assay. ** and *** indicate different from wild-type at the corresponding time at P<0.01 and 0.001, respectively (one-way ANOVA and Dunnett’s test). Horizontal lines in all scatter plots indicate the mean; errors are SEM. n.s. – not significant.

**Fig. S4.**
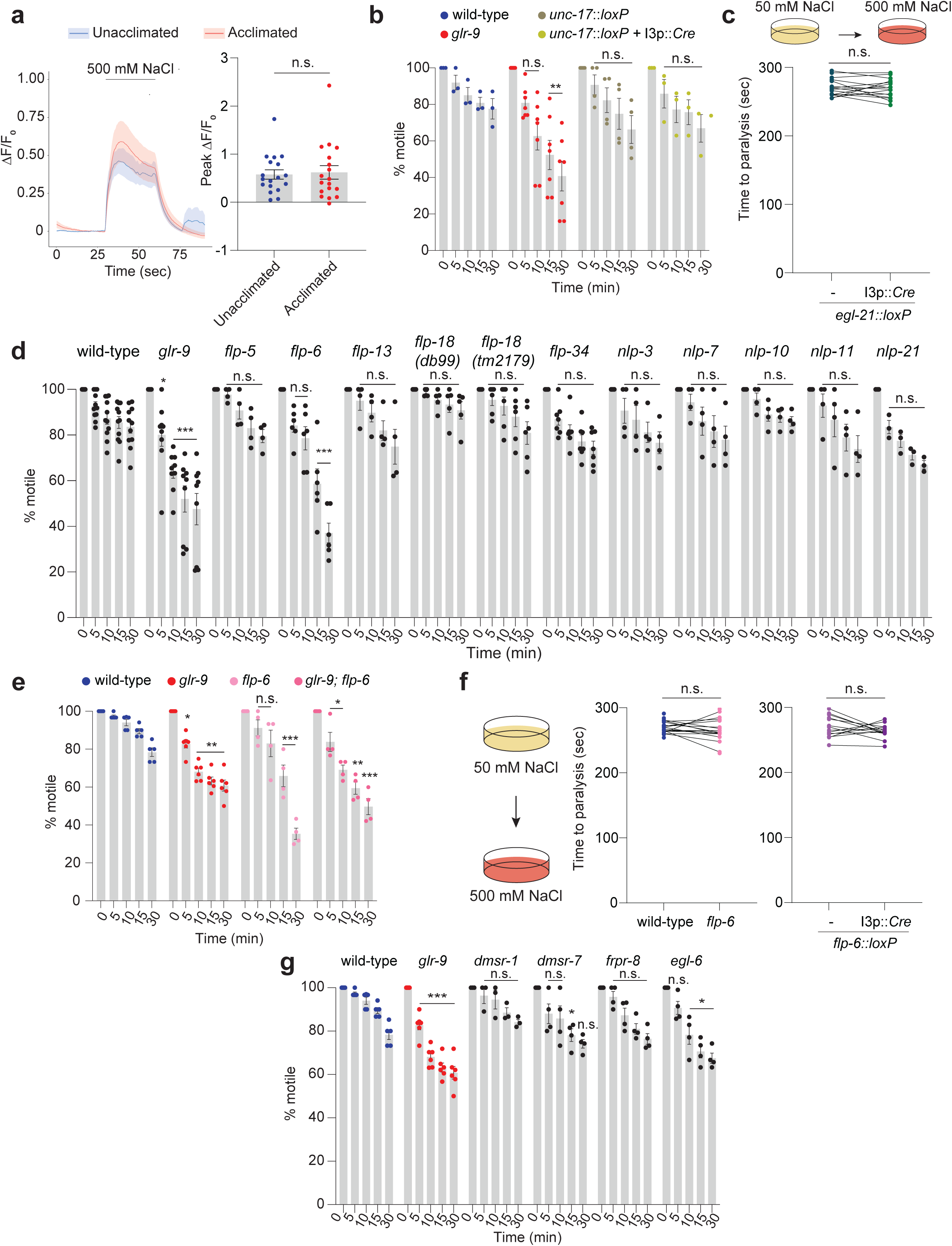
Peptidergic but not cholinergic signaling from I3 is required for salt acclimation. **a)** (Left) Mean GCaMP6s fluorescence change in I3 in response to 500 mM NaCl prior to and following acclimation to 250 mM NaCl for 3 hrs. (Right) Peak fluorescence intensity changes calculated from traces on the left. Each dot is the value from a single I3 neuron. **b,d,e,g)** Percentage of animals of the indicated genotypes that are motile at each time point following a shift from the acclimation plate (250 mM NaCl for 3 hrs) to 500 mM NaCl. Alleles used were *flp-5(tm10075), flp-6(ok3056), flp-13(tm2427)*, *flp-18(db99), flp-18(tm2179), flp-34(ok3071), nlp-3(tm3023), nlp-7(tm2984), nlp-10(tm6232), nlp-11(syb530), nlp-21(tm2569), dmsr-1(sy1522), dmsr-7(sy1539), frpr-8(sy1362)* and *egl-6(n4537)*. Cre was expressed in I3 under the *glr-9* promoter. Each dot is the value from a single assay; n=30 animals per assay. *, ** and *** indicate different from wild-type at the corresponding time at P<0.05, 0.01, and 0.001, respectively (one-way ANOVA and Dunnett’s test). For e and g, wild-type and *glr-9* data were interleaved, and are repeated. **c,f)** Quantification of time to paralysis of animals of the indicated genotypes in the shown assay conditions (cartoons at top (c) or left (f)). Each dot is the time at which all ten animals in a single assay are immotile. Cre was expressed in I3 under the *glr-9* promoter. Control and experimental animals were examined in parallel in the same assay. The *glr-9(oy180)* allele was used in all experiments. Horizontal lines in all scatter plots indicate the mean; errors are SEM. Data shown are from 2-3 independent experiments each. n.s. - not significant.

**Fig. S5.**
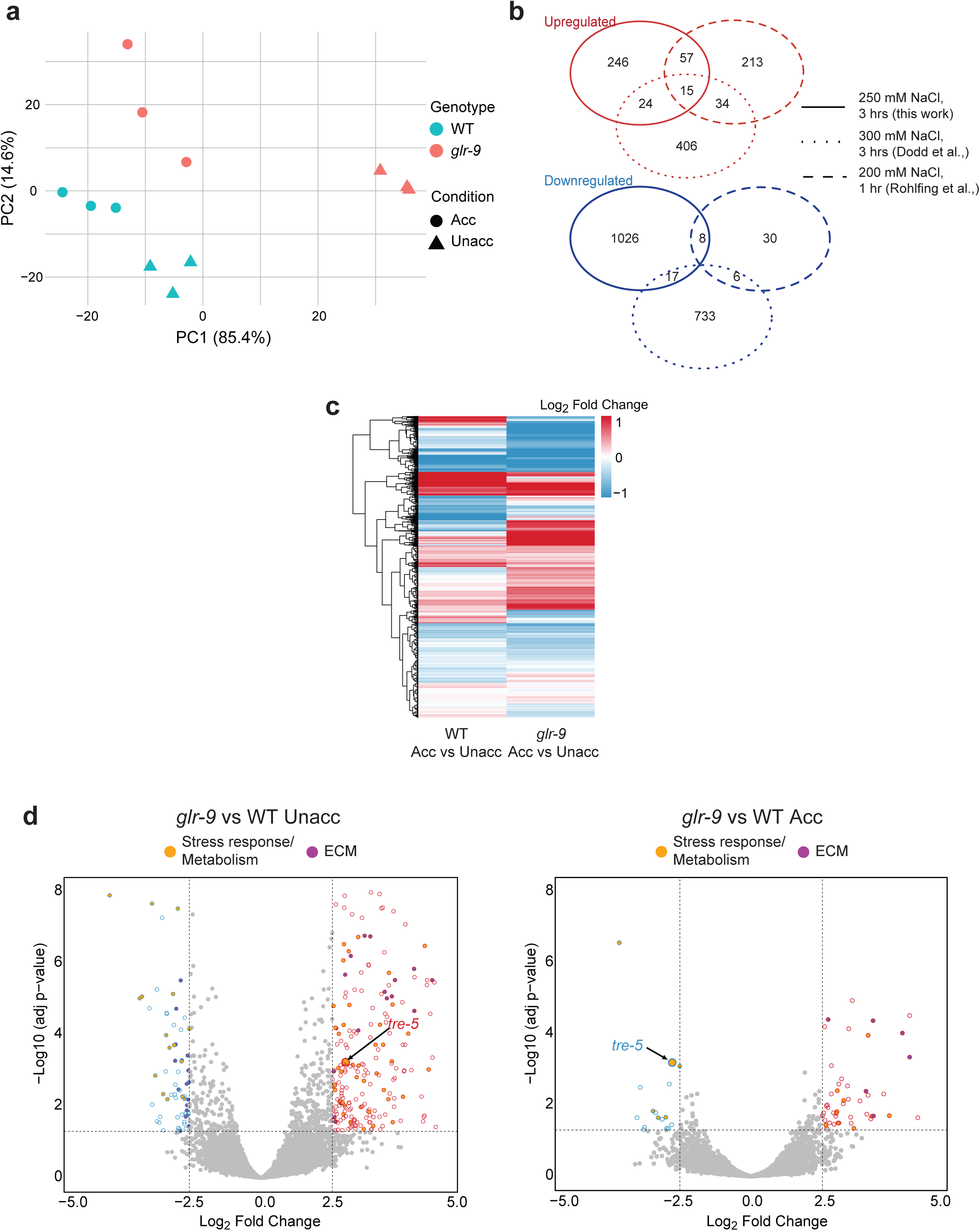
Salt acclimation-induced changes in gene expression are partly regulated by GLR-9-mediated signaling. **a)** Principal component analysis plot of RNA-Seq data clustered by genotype and condition. Each data point is a single replicate. **b)** Venn diagram showing overlap of differentially expressed genes between unacclimated and salt acclimated wild-type animals identified in this and previously published work (Dodd et al., 2018; Rohlfing et al., 2011). **c)** A hierarchically clustered heatmap of all differentially expressed genes between acclimated and unacclimated wild-type and *glr-9* mutants. **d)** Volcano plots of differential gene expression between unacclimated and salt-acclimated wild-type and *glr-9(oy180)* animals. Horizontal and vertical dashed lines indicate significant at adjusted p-value <0.05 and log_2_ fold change >2 or <−2, respectively. A subset of genes involved in osmotolerance and discussed in this work is indicated. Molecules categorized as being involved in stress responses or metabolism, and in extracellular matrix remodeling, are indicated in yellow and purple, respectively.

**Fig. S6.**
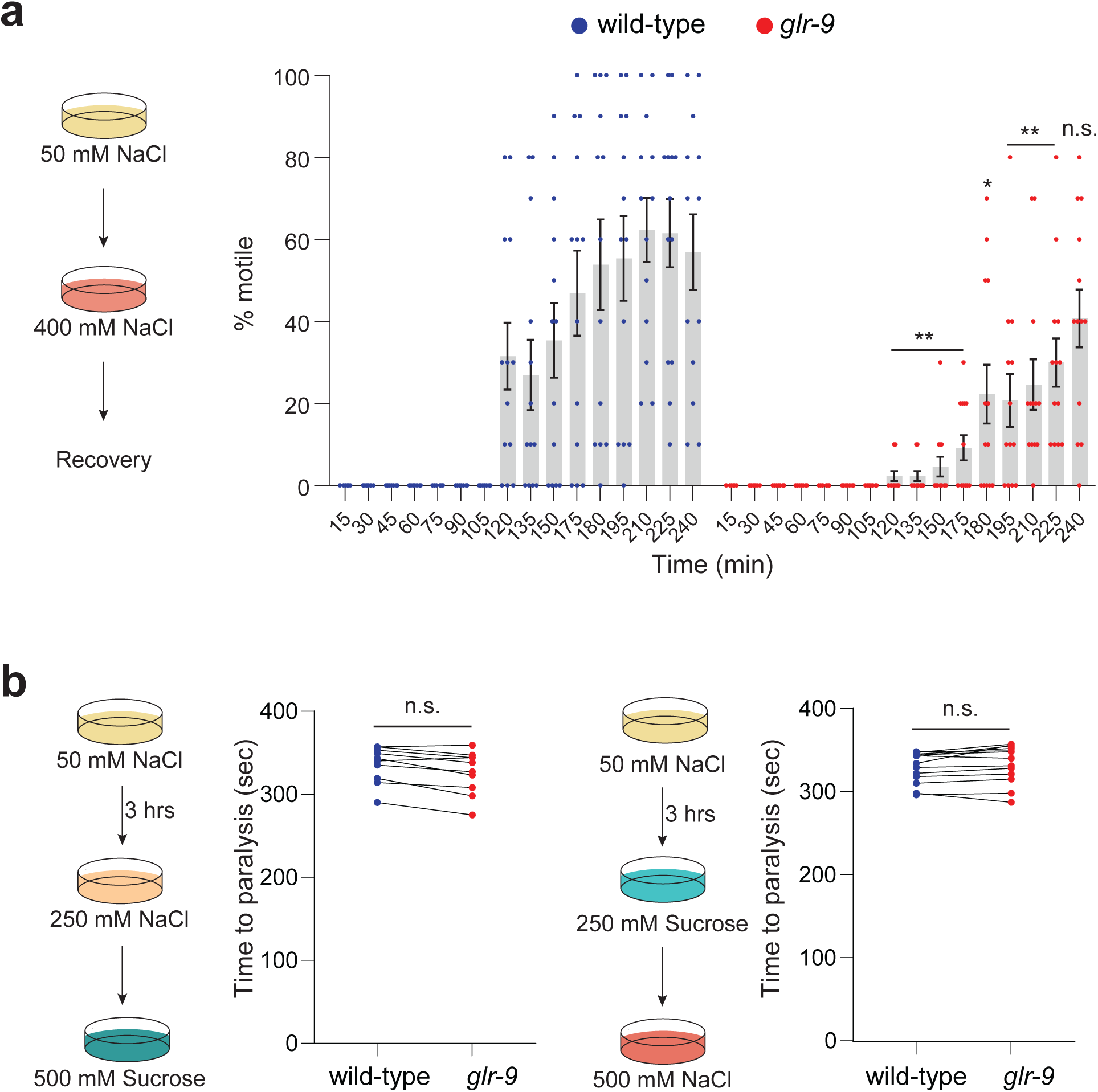
*glr-9* mutants exhibit reduced recovery following salt stress. **a)** Percentage of wild-type or *glr-9(oy180)* mutants that are motile at each time point after being moved from 50 mM NaCl to 400 mM NaCl. Each dot is the value from a single assay; n=10 animals per assay. * and ** indicate different from wild-type at the corresponding time at P<0.05, 0.01, and 0.001, respectively (one-way ANOVA and Dunnett’s test). n.s. – not significant. **b)** Quantification of time to paralysis upon shifting wild-type or *glr-9(oy180)* mutants acclimated to 250 mM NaCl or 250 mM sucrose to 500 mM sucrose or 500 mM NaCl, respectively. Each dot is the time at which all ten animals in a single assay are immotile. Wild-type and *glr-9* mutants were examined in parallel in the same assay. n.s. – not significant.

**Movie S1.** (Right) 3D-rendered image of GLR-9::GFP localization at the I3 sensory ending (Left) generated using Imaris Analysis Software (version 10.2.0).

**Movie S2.** 3D reconstruction of the I3 sensory ending from electron microscope serial sections.

**Data Table 1.** List of upregulated and downregulated genes and gene categories across each condition/genotype from RNA-Seq analysis. The category list is ordered from highest to lowest gene count.

**Table S1.**
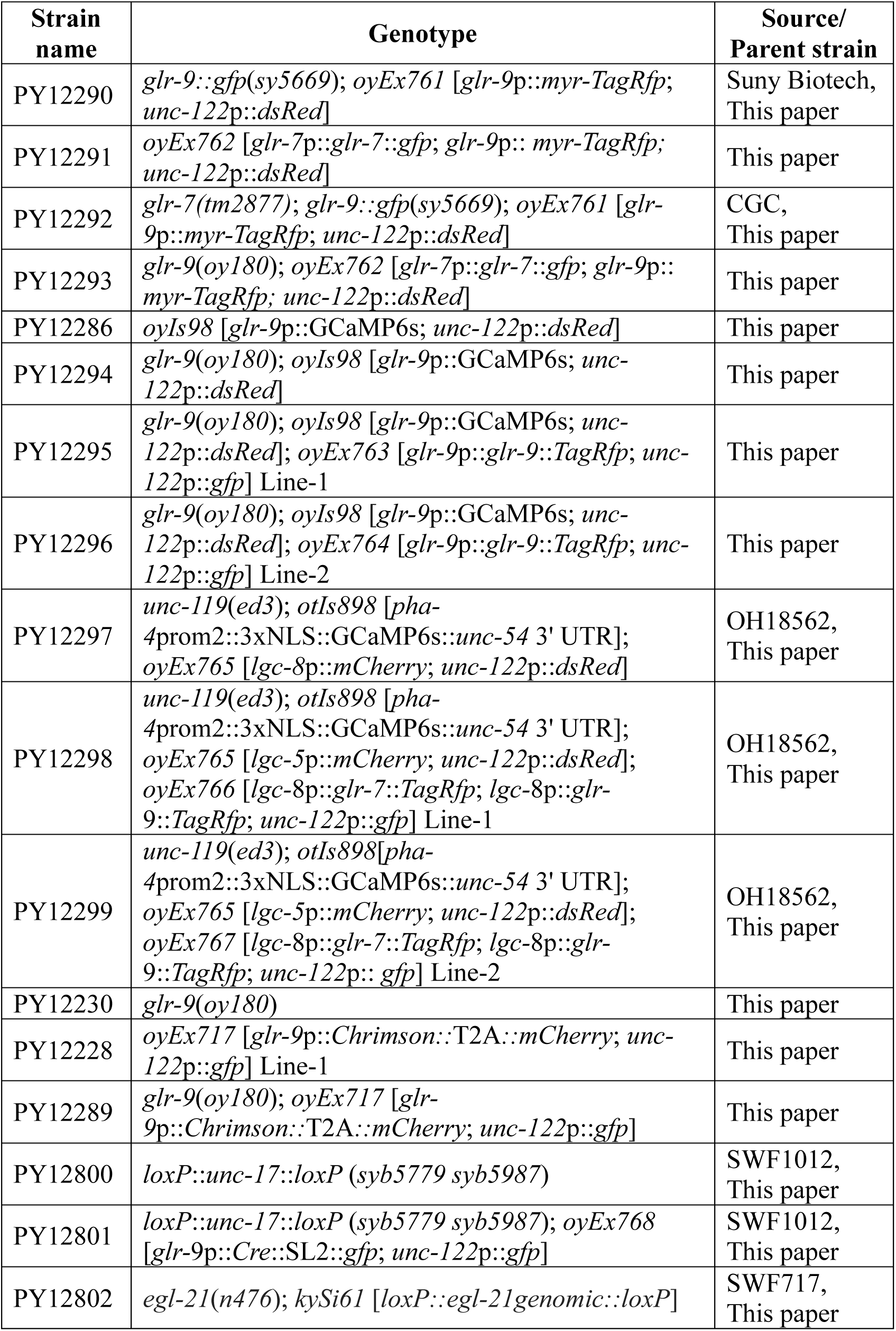
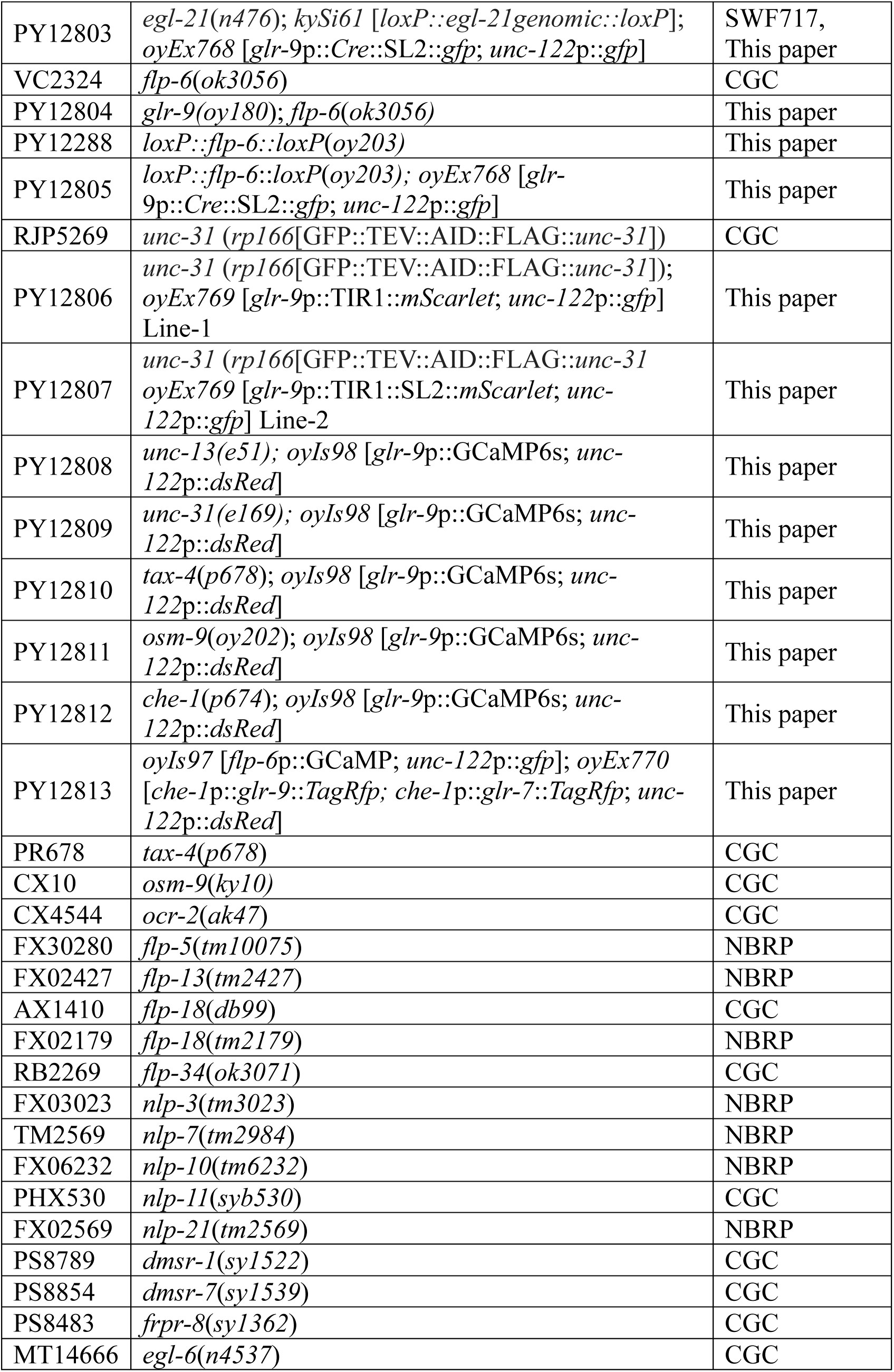
List of strains used in this work.

**Table S2.** List of plasmids used in this work.

| Plasmid name | Description | Source |
| --- | --- | --- |
| PSAB1374 | *glr-9p::myr-TagRfp* | This paper |
| PSAB1373 | *glr-7p::glr-7::gfp* | This paper |
| PSAB1383 | *glr-9p::GCaMP6s* | This paper |
| PSAB1375 | *glr-9p::glr-9::TagRfp* | This paper |
| PSAB1376 | *lgc-8p::mCherry* | This paper |
| PSAB1378 | *lgc-8p::glr-7::TagRfp* | This paper |
| PSAB1377 | *lgc-8p::glr-9::TagRfp* | This paper |
| PSAB1371 | *glr-9p::Chrimson::T2A::mCherry* | This paper |
| PSAB1379 | *glr-9p::Cre::SL2::gfp* | This paper |
| PSAB1380 | *glr-9p::TIR1::mScarlet* | This paper |
| PSAB1381 | *che-1p::glr-7::TagRfp* | This paper |
| PSAB1382 | *che-1p::glr-9::TagRfp* | This paper |

